# Context-dependent spatial multicellular network motifs for single-cell spatial biology

**DOI:** 10.1101/2025.05.05.651422

**Authors:** Amos Zamir, Yael Amitay, Yuval Tamir, Leeat Keren, Assaf Zaritsky

## Abstract

The clinical state of diseased tissue is caused by complex intercellular processes that go beyond pairwise cell-cell interactions and are difficult to infer due to the combinatorial explosion of such high-dimensionality. We present context-dependent identification of spatial motifs (CISM), a two-step method to identify local cell structures associated with a disease state in single cell spatial data. First, for each tissue, CISM enumerates structures of enriched reoccurring multicellular patterns that define modular ‘motifs’ in the multicellular network. Second, discriminative motifs are selected according to the context - their presence in patients at different clinical disease states. By applying CISM, we show that modular structures composed of as little as 3-5 cells and their relative spatial arrangement can encode differences in clinical disease states in cohorts of triple-negative breast cancer (TNBC) and melanoma patients. Machine learning validation indicated that discriminative motifs outperform state-of-the-art methods for disease state prediction while enabling interpretation of which interactions in what spatial context are associated with these predictions. CISM-derived discriminative motifs may define an intermediate spatial scale of abstraction and modularity in multicellular organization and function with broad applicability in the domain of spatial single cell omics and beyond.

## Introduction

Individual cells communicate with and respond to neighboring cells through a complex interplay of chemical and physical cues. These local cell-cell interactions are integrated across space and time to form an emergent tissue organization and structure. In disease, disruption of molecular and cellular processes can lead to abnormal tissue organization and physiological dysfunction. Deciphering the relations between tissue structure and tissue physiology is the holy grail of tissue biology and pathology. Spatial single cell ‘omics’ technologies enable to determine for every cell its cell type and state in its physiological context *in situ*. These technologies define the next technical challenges of systematically representing organization and bridging the different spatial scales from individual cells to physiologically meaningful tissue states (Palla et al., 2022). Spatial analysis efforts are currently mostly focused on the macro spatial scales of the full tissue (Ali et al., 2024; Tamir et al., 2024; Wu et al., 2022) and tissue compartments/macroenvironments (Amitay et al., 2024; Blise et al., 2022; Greenwald et al., 2025; Keren et al., 2018; Risom et al., 2022), and the micro spatial scales of cellular microenvironments/neighborhoods/niches/communities (Ali et al., 2024; Amitay et al., 2024; Birk et al., 2024; Dayao et al., 2023; Jackson et al., 2020; Samorodnitsky et al., 2024; Wu et al., 2022) and pairwise interactions between different cell types (Ali et al., 2024; Dayao et al., 2023; Keren et al., 2018; Risom et al., 2022; Seal et al., 2024), and the intersection (Risom et al., 2022) and the integration between these different spatial and organizational scales (Amitay et al., 2024; Greenwald et al., 2025; Jackson et al., 2020). An open challenge remains in quantifying, and then understanding, intermediate spatial and organizational scales. For example, revealing local multicellular organization principles that extend beyond pairwise interactions and deciphering how local cellular interactions scale up to influence global tissue organization and physiological state.

We hypothesize the existence of an intermediate spatial scale of tissue organization - a modular multicellular unit involving a few locally interacting cells that can define different clinical states of diseased tissues. Such modular components can define a new layer of abstraction that can help reveal biologically meaningful higher-order intercellular interactions that are associated with tissue state, thus bridging the gap between pairwise interactions and tissue scale organization. The main limitation preventing computational analysis of these fine-scale higher-order structures is the number of putative intercellular interactions that grow exponentially in the number of cells and thus are difficult to explicitly infer systematically.

Here, to characterize higher-order fine-scale organization of multicellular spatial networks of human disease tissues, we present a two-step method that first reduces the number of potential higher-order interactions with unsupervised selection of local multicellular structures and then associates these higher-order interactions with the tissue disease state. Specifically, we adapt and extend the concept of “network motifs”, reoccurring sub-networks that define the “building blocks” of complex networks (Milo et al., 2002). Seminal studies established motifs in transcription networks of multiple experimental systems and organisms, suggestive of their role as modular and universal building blocks of transcription networks that are repeatedly selected by evolution for their biological function (Alon, 2007). Motifs also appeared in other biological and non-biological networks (Milo et al., 2004). We introduce a new method named <u>C</u>ontext-dependent Identification of <u>S</u>patial <u>M</u>otifs (CISM) that identifies discriminative motifs in the spatial network that are associated with human disease states. Applied to investigate future metastases in melanoma and in short- versus long-term patient survival in triple-negative breast cancer, we demonstrated that CISM discriminative motifs of 3-5 cells represent a signature for tissue disease state. These discriminative motifs were used as features for accurate machine learning prediction, and their localization in the tissue revealed fine-scale spatial contexts associated with the disease state. Specifically, in metastasis-free sentinel lymph nodes of stage II melanoma patients, CISM identified discriminative motifs consisting of B cells and CD8 T cells that appeared in the outer layers of the germinal centers as a precursor for future remote metastases. In TNBC patients, CISM identified a discriminative motif consisting of interactions between tumor cells, CD4 T cells and dendritic cells that was associated with long-term survival. Overall, CISM revealed an intermediate spatial scale of emergent intercellular organization involving as little as 3-5 cells and their relative spatial composition. These results define a new layer of abstraction and modularity in tissue organization and function associated with disease tissue state. CISM is made publicly available as open source and can be applied to any network representation in the broad domain of spatial biology.

## Results

### <u>C</u>ontext-dependent <u>I</u>dentification of <u>S</u>patial <u>M</u>otifs (CISM)

We introduce context-dependent identification of spatial motifs (CISM) as a bottom-up method for grouping cells into fine-grained spatially resolved multicellular functional modules that provide meaningful information regarding physiological tissue states (Fig. 1). Following single cell segmentation and cell type classification (Fig. 1A), we define a *spatial multicellular network* representing the spatial organization of a tissue, where the nodes of the network are single cells with an edge between every pair of adjacent cells according to the Delaunay triangulation (Fig. 1B, left). This process creates a “colored” graph, where each node is colored according to its corresponding cell type. Network motifs are defined as sub-graphs (i.e., a set of cells and edges) that occur more than expected in the graph, where “more than expected” is determined according to a statistical test. Intuitively, a subgraph is considered a motif if its frequency in the observed graph is significantly larger than its frequency in random graphs generated by switching the edges in a manner that preserves the degree distribution (Methods). Existing methods for motif identification in “colored” graphs are bounded in the number of colors as a function of the motifs’ size (Smoly et al., 2017). This is a substantial limitation for single cell multiplexed data that is characterized by multiple cell types. To address this challenge, we implemented FANMOD+, an adjusted implementation of an existing method called FANMOD (Wernicke & Rasche, 2006), by replacing a 64-bit with a 128-bit subgraph encoding (see Methods for more details). FANMOD+ can support up to 128-colored networks for motifs of size 3-6 cells, going well beyond existing motif detection tools that support at most 15 colors for motifs of size 4-6, 3 colors for motifs of size 7 and 1 color (i.e., no node coloring) for motifs of size 8 (Smoly et al., 2017).

**Figure 1.**
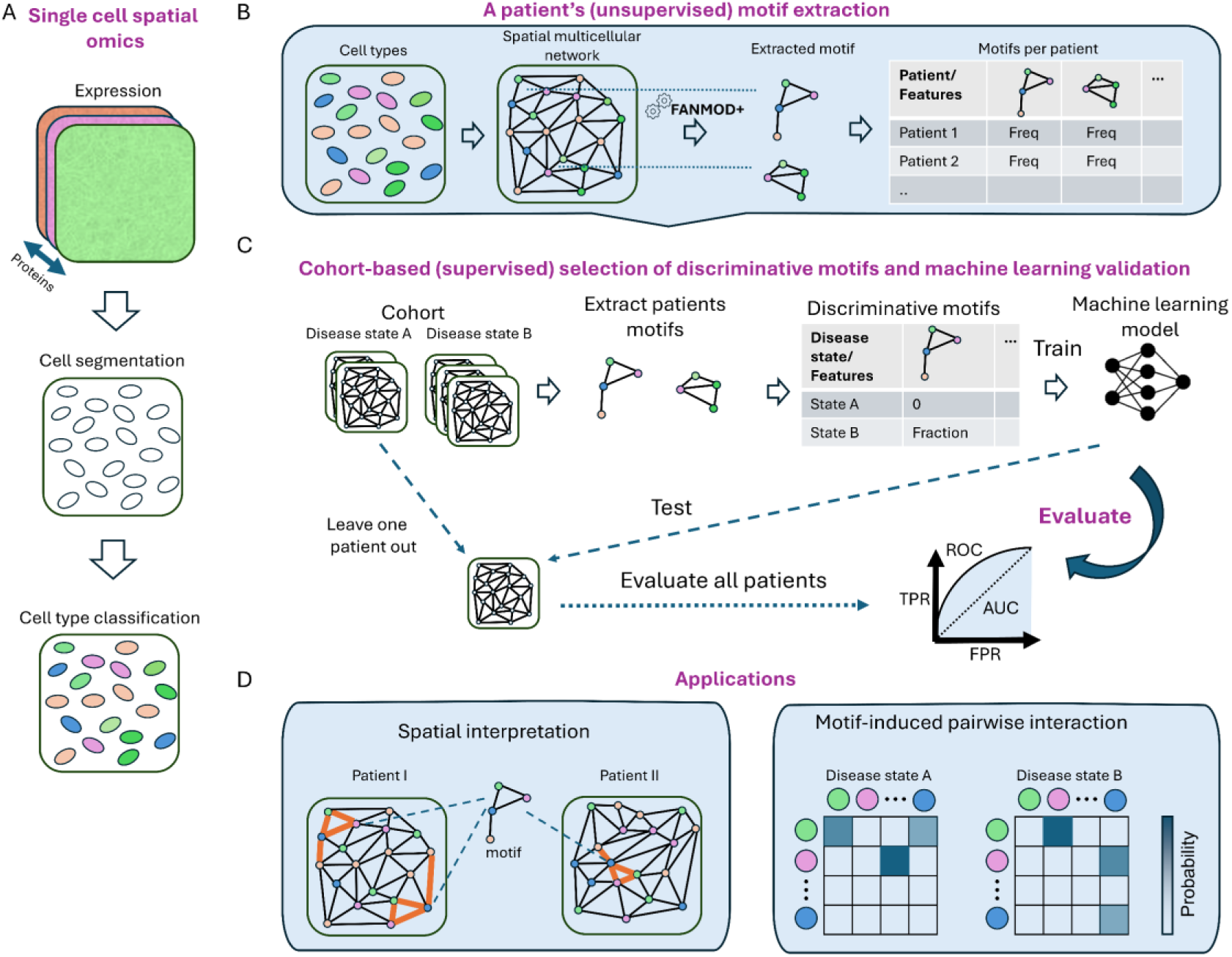
Extraction of spatial motifs, context-based selection, machine-learning evaluation, and visualization. (**A**) Single-cell type classification (top-to-bottom): multi-channel spatial omics data at single-cell resolution, cell segmentation, cell type classification. (**B**) Extract motifs using FANMOD+ from each patient and record the relative frequency of each motif. (**C**) Discriminative motifs and machine learning evaluation. Left-to-right: leave one patient out, extract motifs, select discriminative motifs, train a machine learning model to predict the disease tissue state using discriminative motifs as features, and evaluate the model on the left-out patient. Repeat this process for all patients and calculate the ROC according to the pooled prediction probabilities. (**D**) Applications. (1) Spatial Interpretation (left): moving back to space and interpreting. Overlaying the motifs on the spatial multicellular network – the motifs’ localization in the tissue can infer the spatial contexts associating a motif to a disease state. (2) Exploratory analysis of motif-induced pairwise cell-cell interactions (right): the probability of an edge between two cell types in the pooled set of discriminative motifs instances can define a pairwise interactions-based disease state signature.

Given a motif size *n* (i.e., number of cells), for each patient’s tissue sample, CISM extracts and enumerates all the motifs from the huge landscape of all subgraphs and calculates for each motif its frequency with respect to the total number of corresponding subgraphs of size *n* in the patient’s sample (Fig. 1B, right). To identify motifs that are associated with a clinical parameter such as disease state, CISM selects the motifs that appear in at least a predetermined fraction of the patients with the corresponding disease state, and that do not appear in any patients with other disease states; we term these as *context-dependent discriminative motifs* (onwards referred as discriminative motifs). These discriminative motifs can provide biologically-meaningful interpretable information, at the scale of a few cells and their putative local interactions. Altogether, the motifs are extracted in an unsupervised fashion, and then selected, with supervision, according to a discrimination criterion, such as disease state or clinical treatment.

CISM context-dependent discriminative motifs can be used to formulate a machine learning-based prediction of the tissue’s disease state. Each patient is represented by the (sparse) vector of its discriminative motifs’ frequencies. Since patient cohorts are commonly limited to dozens of patients, CISM evaluates the discrimination capacity of this motif’s representation in leave-one-patient-out cross validation: multiple rounds of training and testing, each round with one patient designated for a test, and all other patients are used as the training set. More specifically, each iteration followed the processes depicted in Fig. 1C (left-to-right): (1) keep one patient out, (2) extract discriminative motifs from all patients excluding the one that was left out, (3) use the most frequent discriminative motifs as representations to train a machine learning model to predict the tissue disease state, (4) apply the model on the held-out patient. Importantly, in this setting, the training process is fully independent of the patient in the test (Jones, 2019). This process was repeated for all patients, and the different models’ prediction probabilities of each held-out patient were pooled and evaluated with the area under the receiver operating characteristic curve (AUC-ROC). Discriminative motifs can be evaluated based on their prevalence across patients, their contribution to the models’ performance, identity of the cells that comprise them and their pairwise interactions and their spatial location in respect to more global tissue compartments for cross-scale biological interpretation (Fig. 1D).

### CISM classification of melanoma disease state

We first tested CISM on a dataset of multiplexed imaging of sentinel lymph nodes of melanoma patients (Amitay et al., 2024). The cohort consisted of sentinel lymph nodes of 38 stage I-II melanoma patients, who were followed up for at least five years, and classified according to the development of metastases in distant organs (Fig. 2A). A patient with tumor-free lymph nodes and no distant metastases in five years was defined as negative-negative (NN). Correspondingly, those patients with remote metastases five years following diagnosis were defined as negative-positive (NP). The dataset included 76 multiplexed images acquired by MIBI-TOF (Keren et al., 2019), each one containing images of 39 proteins *in situ*. We used the cell type classification (Methods) to assign each cell to one of 15 lymph node microenvironment cell types according to their lineage relationships (Fig. S1, Methods), and defined the corresponding spatial multicellular network according to cell-cell spatial adjacency (Methods). The cell type distribution, defined by pooling all cells from all patients, showed some differences between NN and NP patients with a higher fraction of CD4 T-cells and Macrophages in NN and Germinal Center cells, and Memory CD4 T-cells in NP (Fig. 2B), in accordance with (Amitay et al., 2024). The distribution of pairwise interactions according to the cell-cell edge probabilities showed enrichment of interactions between adjacent pairs of Memory CD4 T cells and between adjacent pairs of B cells (Fig. S2). Notably, Memory CD4 T cells and B cells were the two cell types most abundant in the cell type distribution and appeared for both NN and NP patients. These representations of cell type distribution and pairwise interaction distribution were insufficient for robust discrimination with Random Forest machine learning classification AUC of 0.6 and 0.56 correspondingly (see Methods).

**Figure 2.**
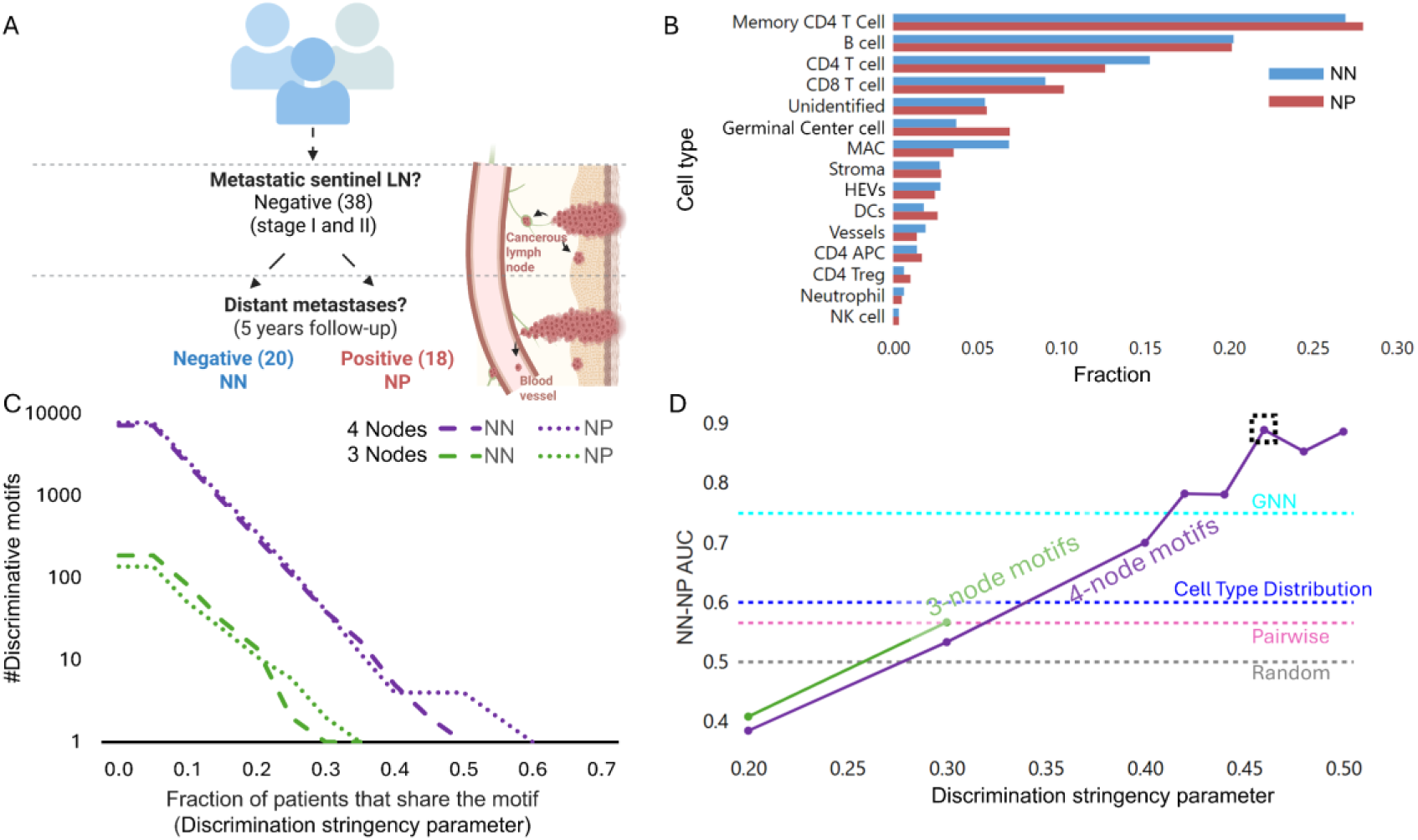
Context-dependent identification of spatial motifs (CISM) can predict melanoma disease state. (**A**) The melanoma cohort includes 38 patients diagnosed with a primary tumor with no apparent tumor cells spreading to the lymph node (i.e., melanoma stage I-II). 20 of these patients did not develop distant metastases (Negative-Negative, NN), while the remaining 18 did (Negative-Positive, NP). (**B**) Cell type distribution for NN and NP patients. (**C**) The number of 3-cell (green) and 4-cell (purple) context-dependent discriminative motifs (y-axis, log scale) over the discrimination stringency parameter (x-axis) - a fraction of patients of a given disease state that share a discriminating motif when none of the patients with the other disease state share it. The dashed line and the dots represent the number of discriminative motifs in NN and NP correspondingly. (**D**) Machine learning validation of melanoma disease state discrimination with CISM’s discriminative motifs representation. Leave one patient out cross-validation (LOOCV) model performance (AUC) (y-axis) over the discrimination stringency parameter (x-axis) (green - 3-cell motifs, purple-4-cell motifs). The pink, cyan, and black dashed horizontal lines represent the LOOCV AUC score of pairwise, GNN, and a random null model, correspondingly. The discriminative motifs derived from 4-cell motifs with the discrimination stringency parameter of 0.46 for discriminative motifs selection were used for further analysis (see Methods for parameter setting justification). The sensitivity to the discrimination stringency parameter and to the motif’s size is shown in Fig. S8 and Fig. S9 correspondingly.

We applied FANMOD+ to extract hundreds-of-thousands to thousands of 3-, 4-, and 5-cell motifs per patient, correspondingly, where the majority of these motifs were deemed highly statistically significant with p-value < 0.002 (Fig. S3A). To examine the composition of these motifs, we computed the cell type distribution and the pairwise cell-cell interactions distribution across the pooled set of 4-cell node motif instances in all patient samples (Fig. S3B-C). Both the motifs-induced cell type distribution and the motifs-induced pairwise interactions were similar to the corresponding distribution in the general distribution, before motifs extraction (compare Fig. S3B-C to Fig. 2B and Fig. S2 correspondingly) and did not show differences between NN and NP patients. This step reduced the number of subgraph instances by two-orders of magnitude (Table S1). The exclusion of non-motif subgraphs distills the space of subgraphs from those subgraphs that may appear by chance, thus enabling the second step of identifying “discriminative” motifs that exclusively appear in patients of one disease state.

Next, we performed context-dependent motif selection according to the discrimination stringency parameter, which is the fraction of patients of a given disease state (i.e., NN versus NP) that share a discriminating motif. This analysis showed similar numbers of discriminative motifs in NN and NP that exponentially decayed as a function of the fraction of the patients with the corresponding disease state that shared the motif (Fig. 2C). To systematically test whether discriminative motifs can provide a rich representation of the tissue disease state, we turned to machine learning validation. Specifically, we performed Random Forest-based leave-one-patient-out cross-validation where each patient was represented by a (sparse) feature vector of its discriminative motifs frequencies. Gradually increasing the discrimination stringency parameter revealed a trend of enhanced classification performance for discriminative motifs that appear in more patients for 3- and 4-cell motifs (Fig. 2D). The classification reached a maximal AUC score of 0.88 with 4-nodes discriminative motifs that appeared (in the training) in at least 46% of patients from one disease state and did not appear in patients from the other disease state. Setting the discrimination stringency parameter to this value had an additional benefit in limiting the overall number of discriminative motifs for downstream interpretation to 19 distinct motifs: 8 that were associated with NN (Fig. S4), and 11 discriminative motifs associated with NP (Fig. S5). Motifs of size three led to inferior classification performance due to a lack of sufficient numbers of discriminative motifs (Fig. 2C-D), and 5-cell motifs reached an AUC score of 0.81 (discrimination stringency parameter = 0.2) that fluctuated and then dramatically dropped for stringent discrimination criteria beyond 0.46 due to training overfitting with few motifs per leave-one-out iteration (Fig. S6). Random forest disease state classification based on the 4-cell discriminative motifs representation surpassed the simple representations of pairwise cell adjacencies and of cell type distributions that reached an AUC of 0.56 (std = 0.02, Fig. 2D – pink horizontal line) and AUC of 0.6 (std = 0.03, Fig. 2D – blue horizontal line), correspondingly (see Methods for full details). The state-of-the-art alternative classification approache of whole graph neural networks (GNN) (Tamir et al., 2024) reached an AUC of 0.7493 (Fig. 2D – cyan horizontal line). To confirm that these results were not due to overfitting, we permuted the patients’ disease state labels and performed the full CISM classification pipeline of leave-one-patient-out cross-validation, including independent identification of discriminative motifs in every round. Repeating this permutation test 100 times produced AUC scores that were distributed around random (mean = 0.4912, std = 0.2138, p-value = 0.01, Fig. S7). Cumulatively, CISM showed remarkable clinical outcome prediction for a hard classification problem that could not be solved with state-of-art approaches. These results indicate that discriminative motifs can serve as multicellular modular signatures of a disease state with the fine spatial resolution of four cells and their spatial adjacencies.

### Exploring of the discriminative motifs-induced cell distribution and motifs-induced pairwise cell-cell interactions

To explore the landscape of the nineteen 4-node discriminative motifs selected over the cross-validation iterations of the machine learning analysis (Fig. S4-5) we visualized their cell type compositions (Fig. 3A). We also calculated their induced cell type distribution across the pooled set of instances of discriminative motifs associated with NN or NP across the patient samples (Fig. 3B). Intriguingly, the discriminative motifs-induced cell type distribution was very different from the general cell type distribution (compare Fig. 3B to Fig. 2B). First, cell types that were common in the general distribution were less common in the discriminative motifs-induced distribution. For example, Memory CD4 T cells, that composed over 25% of the general cell type distribution in both NN and NP patients, appeared in less than 10% of the NN-associated discriminative motifs-induced cell type distribution and did not appear at all in NP-associated motifs. Second, cell types that were less prevalent in the general distribution, were more abundant in the discriminative motifs-induced distribution. For example, macrophages and CD8 T cells, that appeared in ∼10% of the cohort’s general cell population, accounted for 16-29% of the NN- and NP-associated discriminative motifs-induced cell type distributions. Third, while the general cell type distributions showed marginal differences between NN and NP patients, the motifs-induced distribution showed dramatic differences. For example, CD8 T cells and B cells were almost twice as probable and five-times as probable, correspondingly, to participate in NP-associated discriminative motifs, while CD4 T cells appeared almost 4 fold more in NN patients. Also, each of the Memory CD4 T cells, Neutrophils and NK cells, exclusively appeared in ∼10% of the NN-associated motifs cell type distribution.

**Figure 3.**
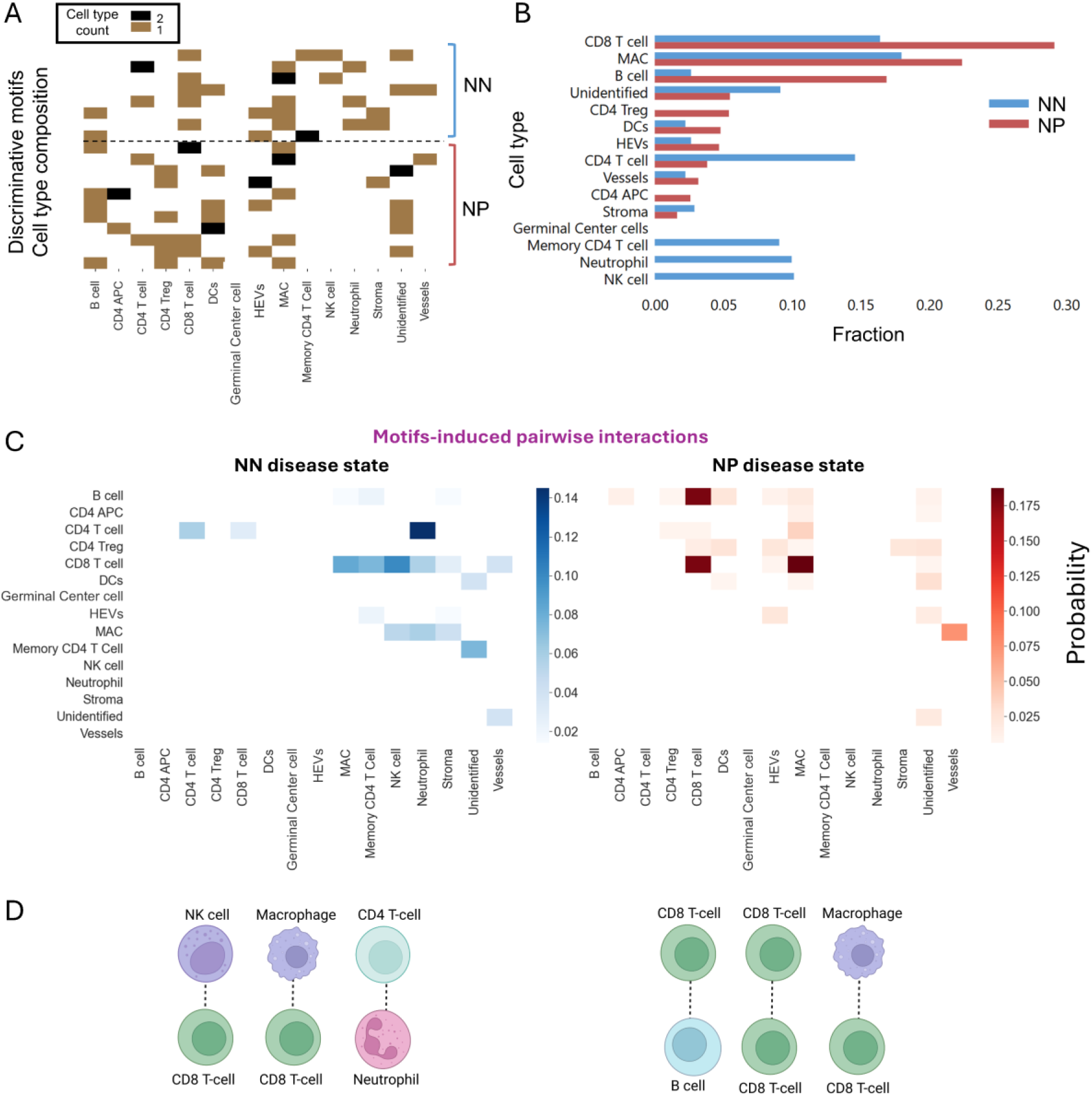
Context-dependent motifs-induced pairwise interactions melanoma disease state. (**A**) The cell type composition of the discriminative motifs associated with NN (n = 8 motifs) and NP (n = 11 motifs) patients. (**B**) The cell type distribution induced from the pooled set of discrmnative motifs’ instances across the cohort. (**C**) The distribution of motifs-induced pairwise interactions. The cell type pairwise edge probabilities in the cohorts’ patients pooled set of instances of discriminative motifs associated with NN (left) and NP (right) patients. Spearman correlation between NN and NP pairwise interactions was -0.0157 with p-value ≈ 0.87. (**D**) The top three highly ranked motifs-induced pairwise interaction.

Next, we examined the distribution of discriminative-motifs induced pairwise cell-cell interactions according to the cell type pairwise edge probabilities in the pooled set of instances of discriminative motifs associated with NN or NP across the patient samples (Fig. 3C). This analysis revealed potential pairwise interactions associated with one disease state and not the other. The three most abundant (motif-induced) pairwise interactions in NN and in NP are depicted in Fig. 3D: Neutrophils with CD4 T-cells, Macrophages with CD8 T-cells, and CD8 T-cells with NK cells, versus B cells with CD8 T-cells, CD8 T-cells with CD8 T-cells, and Macrophages with CD8 T-cells in NN- versus NP-associated discriminative motifs. Intriguingly, the interaction between CD8 T-cells and Macrophages appeared in both NN- and NP-associated discriminative motifs, in different contexts: the (n = 3) corresponding NN-motifs included interactions between CD8 T Cells with Neutrophil, NK cells, or Stroma cells (Fig. S4) while the (n =2) corresponding NP-motifs included interactions between CD8 T Cells with CD8 or CD4 T cells (Fig. S5). The motifs-induced disease state-associated pairwise interactions were robust to the stringency of the motifs’ discrimination criterion (Fig. S8) and to the motif’s size (Fig. S9). These results suggest that the strict context-dependent discriminative motif selection refines the huge and noisy landscape of cell types and all possible cell-cell interactions in the full spatial multicellular network.

### The spatial arrangement of the cells within the motif contributes to the classification of melanoma disease state

A motif is defined according to its cell types and the edges between them. We next asked whether the discriminative motifs’ cell type composition, i.e., the cell types without considering their relative spatial arrangement to one another, is sufficient to discriminate between the NN and NP disease states, or in other words – whether the spatial arrangement of cell types within discriminative motifs matters for disease state classification? To answer this question, we repeated the steps of context-dependent motifs’ selection and machine learning analysis, but instead of using the discriminative motifs we used the discriminative *cell type composition* representations. The cell type composition of a motif was represented as a sparse vector that encodes the number of occurrences of each cell type in the motif and was shown earlier in Fig. 3A. For example, a 4-cell motif that includes two edges from a HEV to two memory CD4 T-cells and two edges from a B-cell to the same two memory CD4 T-cells is transformed to a sparse vector encoding a single B-cell, one High Endothelial Venule (HEV) cell, and two memory CD4 T-cells (Fig. 4A). This cell type composition representation induced fewer discriminative cell type compositions and correspondingly lower values for the discrimination stringency parameter, for example, 21 versus 79 in Fig. S10 versus Fig. 2C for discrimination stringency parameter of 0.3. Thus, preventing us from using the cell type composition representation to evaluate the discrimination. These results indicate that simply using combinatorial interactions of four cells was not sufficient to identify the motifs revealed by CISM and that inclusion of the intra-motif spatial rearrangement contributes to disease state prediction.

**Figure 4.**
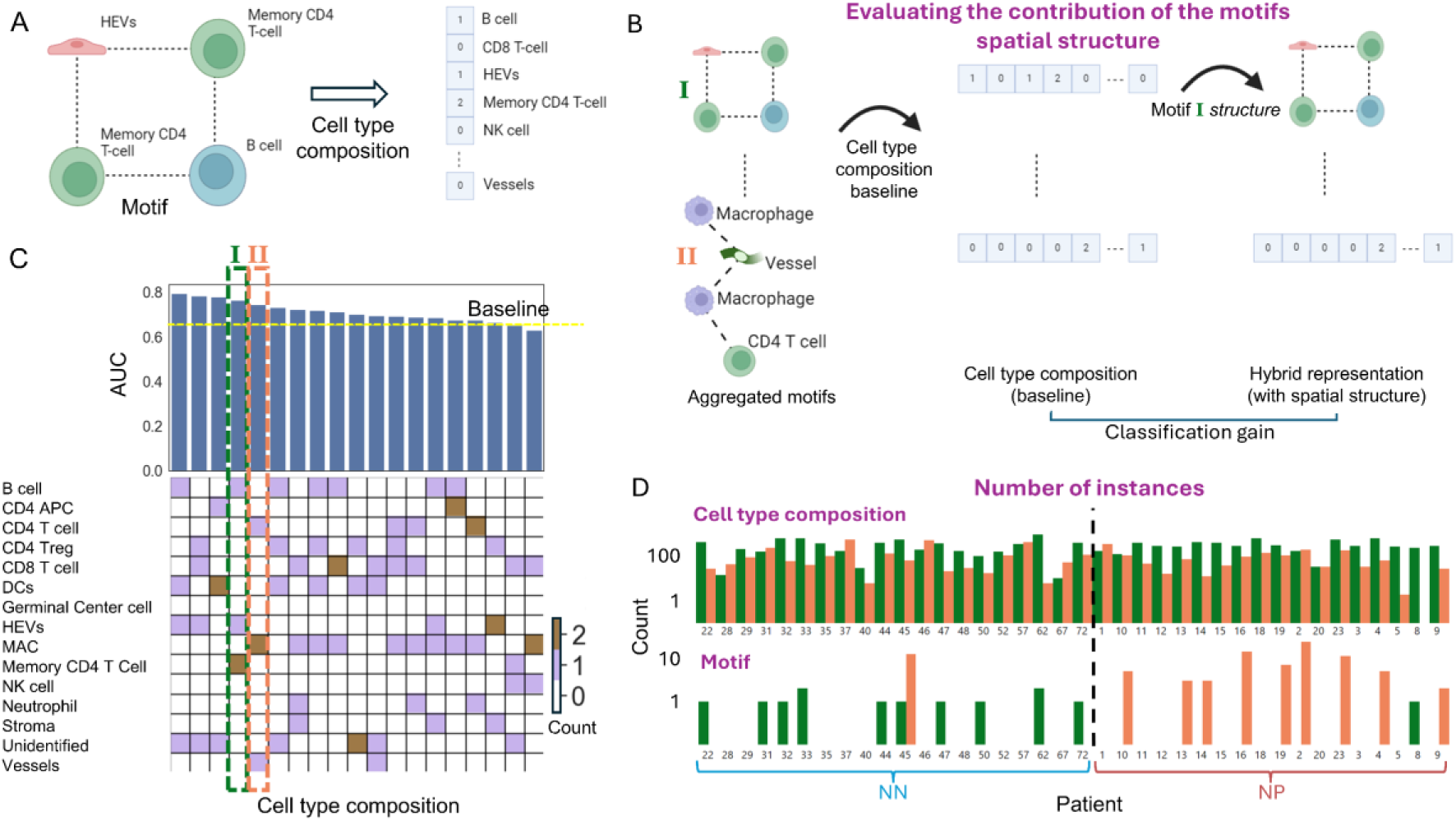
The spatial arrangement of discriminative motifs contributes to disease state classification. (**A**) An example of mapping a discriminative motif (left) to its corresponding cell type composition (right). The cell type composition representation is 15-dimensional, counting the number of instances of each cell type in the input discriminative motif. (**B**) Assessing the discriminative contribution of the spatial arrangement of discriminative motifs. First, all the discriminative motifs (left) are mapped to their corresponding cell type compositions (middle). This discriminative motifs-derived cell type composition representation is used to train a baseline machine learning model to predict the disease state. Second, the contribution of the spatial arrangement is measured as the classification performance gain by replacing one cell type composition with its corresponding discriminative motifs, one at a time (right). (**C**) The ranked classification gain for each cell type composition of the n = 19 discriminative motifs (x-axis). Specifically, each column represents a cell type composition, each row represents a cell type, and the color encodes the number of cell type instances in the corresponding cell type composition. For example, the cell type composition #I corresponds to one B cell, one HEV, and two memory CD4 T cells (see panel A). The deviation of the AUC of a cell type composition baseline (yellow dashed line, AUC = 0.655) with respect to its corresponding hybrid representation (blue bar) is the classification gain. The spatial arrangement of 17/19 of the motifs contributed to the disease state prediction. Two motifs and their corresponding cell type compositions (I – green, II - orange), whose spatial arrangement contributed the most while not containing (less interpretable) “unidentified” cell types, were highlighted for further analysis. (**D**) The number of instances of the cell type compositions I (green) and II (orange) are shown in panels B-C (top) and their corresponding motifs (bottom). The outlier motifs II (patient #45) and I (patient #8) were selected when the corresponding patient was used for the test (i.e., was selected as discriminative when not considering that patient). The cell type compositions did not discriminate between NN and NP (Mann-Whitney U test: cell type compositions I (green): p-value = 0.6398, cell type compositions II (orange): p-value = 0.5586), while the motifs significantly discriminate between NN and NP (motifs I (green): p-value = 0.0029, motifs II (orange): p-value = 0.0032)

To directly assess which of the discriminative motifs’ spatial arrangement are most associated with the disease state, we devised an approach that measures the gain in classification performance attributed to the inclusion of each discriminative motif’s spatial arrangement instead of its corresponding cell type composition. First, we transformed the representation from the discriminative motifs space to the cell type composition space by replacing each discriminative motif with its corresponding cell type composition (Fig. 4B). Note that multiple discriminative motifs can be mapped to the same cell type composition, and that a cell type composition can include instances of motifs that did not meet the strict discrimination stringency parameter used to determine the discriminative motifs. This discriminative motifs-derived cell type composition representation reached a reduced AUC score of 0.655, which was used as a baseline (Fig. 4B, middle). Second, beginning from this discriminative motifs-derived cell type composition representation, we iteratively introduced back the spatial information for each cell type composition by replacing it with the discriminative motifs that induced it. This replacement created a hybrid representation that contained both cell type composition and discriminative motif features (Fig. 4B, right). I.e., each patient was represented by a vector encoding the frequencies of the current set of discriminative motifs and cell type compositions. This hybrid representation was used for machine learning assessment, where the contribution of the corresponding motifs’ spatial arrangement was attributed to the classification gain with respect to the cell type composition baseline (Fig. 4B, “Classification gain” = right - middle). The classification gain for each cell type composition was ranked, showing that the vast majority (17/19) of the motifs’ spatial arrangement contributed to disease state prediction (Fig. 4C). The three highest ranked cell type compositions included “unidentified” cell types, i.e., cells without a specific classification, and thus were not further analyzed (see Discussion). The 4^th^ highest ranked cell type composition contributed 0.11 to the cell type composition baseline AUC, and was composed of one B cell, one high-endothelial venule (HEVs) cell, and two memory CD4 T cells (Fig. 4C, motif #I in green, this motif was used as an example in Fig. 4A-B). The 5^th^ highest ranked cell type composition contributed 0.09 to the baseline AUC and was composed of a CD4 T cell, a pair of Macrophage cells, and a vessel cell (Fig. 4B, motif #II in orange; this motif was used as an example in Fig. 4B).

To further verify that the spatial arrangement of these motifs provides more discriminative power than their corresponding cell type compositions, we counted the number of instances of these motifs and their corresponding cell type compositions across the cohort’s patients. While the number of instances of the cell type compositions induced by motifs I and II did not show clear discrimination between the disease states (Fig. 4D, top), the prevalence of motifs I and II clearly discriminated between NN and NP (Fig. 4D, bottom). Thus, distilling the contribution of the discriminative motifs’ spatial arrangement indicates that the specific intra-motif direct cell-cell interactions are more sensitive as markers for disease state with respect to the corresponding motif’s cell type composition.

### Spatial interpretation of an NP-associated motif localization reveals a “connector” between the B cells surrounding the germinal center and the rest of the immune microenvironment

One of the most appealing attributes of CISM is the ability to locate instances of a discriminative motif in the multicellular network and characterize the stereotypic spatial contexts associated with its localization and the disease state. A discriminative motif consistently localized in a specific microenvironment can drive the interpretation of potential mechanisms regarding the motif’s physiological roles in the context of the disease. To decide which discriminative motifs to spatially interpret, we balanced between two desired properties: (1) generalization – discriminative motifs that appeared in many patients, and (2) interpretability – discriminative motifs with many instances per patient. Placing the 19 NN-NP discriminative motifs on this generalization-interpretability axes highlighted three motifs that appeared in many patients with many instances on average per patient (Fig. 5A-C). Motif I (green) was NN-associated and included a Macrophage, Neutrophil, and a pair of CD4 T cells, motif II (magenta) was also NN-associated and comprised of an NK cell, a CD8 T cell, a Memory CD4 T cell, and an Unidentified cell, and motif III (yellow) was NP-associated and comprised of a Macrophage, a pair of CD8 T cells, and a B cell. Motif III was found to be the most abundant in several NP patients and was thus used for spatial interpretation. Localizing motif III in the tissue revealed a stereotypical spatial pattern specifically appearing at the edges of B cell follicles that surround the germinal center in 7/18 of the NP patients (Fig. 5D, Fig. S11). The motif’s B cell was part of the germinal center-associated B cell follicle that was adjacent to two CD8 T cells that were adjacent to a macrophage. This localization pattern may resemble a “connector” between the germinal center to the rest of the immune microenvironment. Enriched in NP patients, this motif associated with long-term metastatic progression. These results highlighted the possibility of providing hypothesized physiologically meaningful interpretation by observing the spatial localization of CISM-derived discriminative motifs.

**Figure 5.**
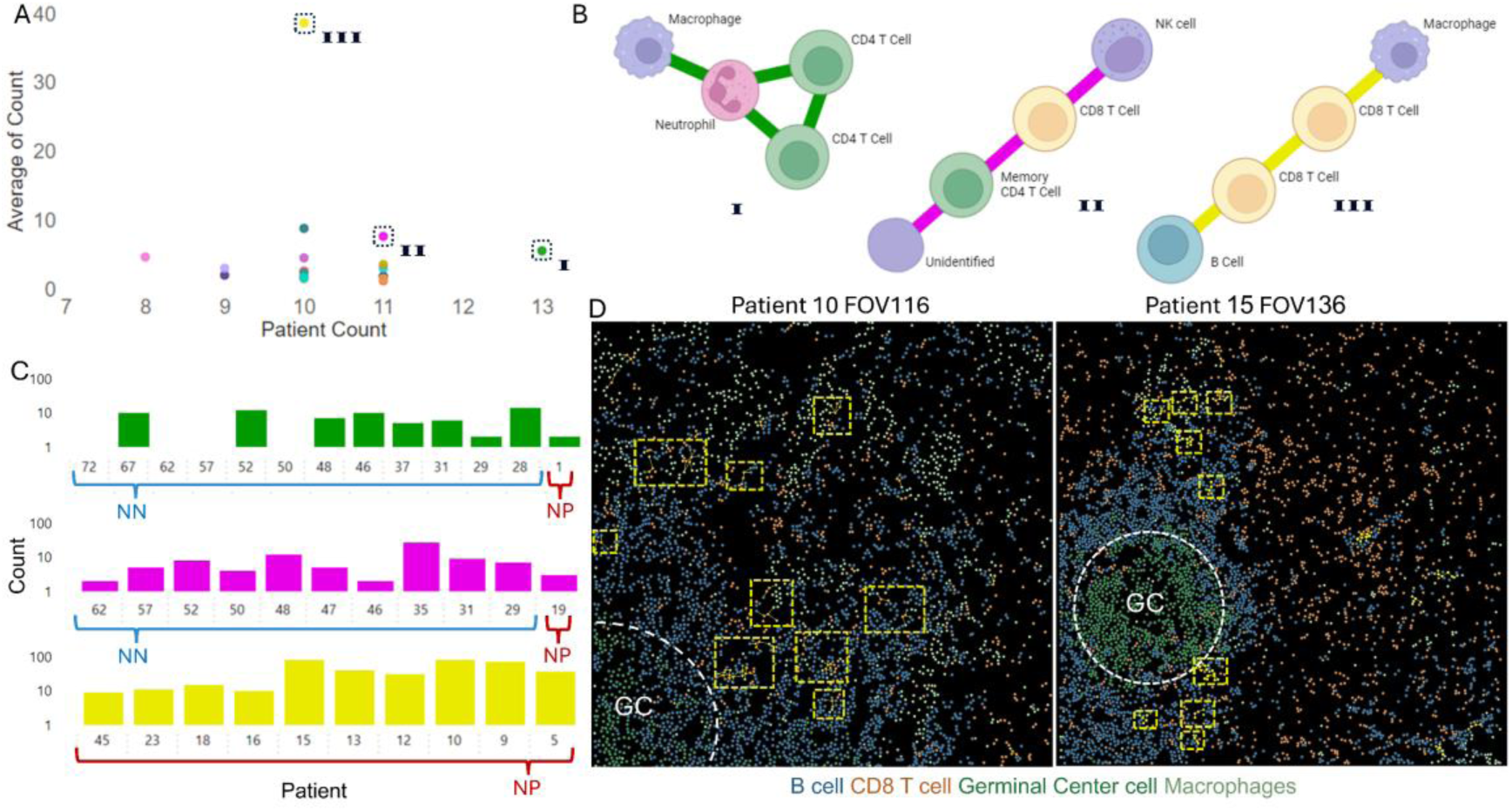
Moving back to space to characterize the microenvironment of abundant discriminative motifs. (**A**) The mean per-patient number of discriminative motif instances (y-axis) in relation to the number of patients that contained that motif (x-axis). Each dot represents a discriminative motif (n = 19). Note motifs I (green), II (magenta), and III (yellow) that appear in many patients and in many instances per patient. (**B**) The three discriminative motifs that were marked in A. The edge colors are consistent across the other figure panels. (**C**) The number of instances of motifs I-III (top-bottom, log scale) across patients. Only patients with at least one instance were listed. (**D**) Localizing motif III in the tissue of NP patients #10 (left) and #15 (right) and in additional patients (Fig. S11). Each dot represents a cell colored according to its cell type: B cell, CD8 T cell, Germinal Center cells, and macrophages (other cell types are not shown). The germinal center (GC) is surrounded by a dashed white circle. Instances of motif III are marked with dashed yellow rectangles (larger rectangles indicate multiple instances of the motif).

### Context-dependent NP-associated discriminative motifs

CISM’s context dependency implies that the same motifs extracted from a specific tissue can be differentially analyzed according to the question at hand – via context-dependent selection of the discriminative motifs. To test how context can alter the motifs-induced pairwise interactions and the spatial interpretation of the motifs’ localization, we turned to the extended melanoma cohort that also included 13 patients that had metastatic sentinel lymph nodes that did not develop distant metastases within a follow-up of at least 5 years (PN) (Amitay et al., 2024). We then evaluated the motifs that discriminated between NP patients and PN patients (Fig. 6A).

**Figure 6.**
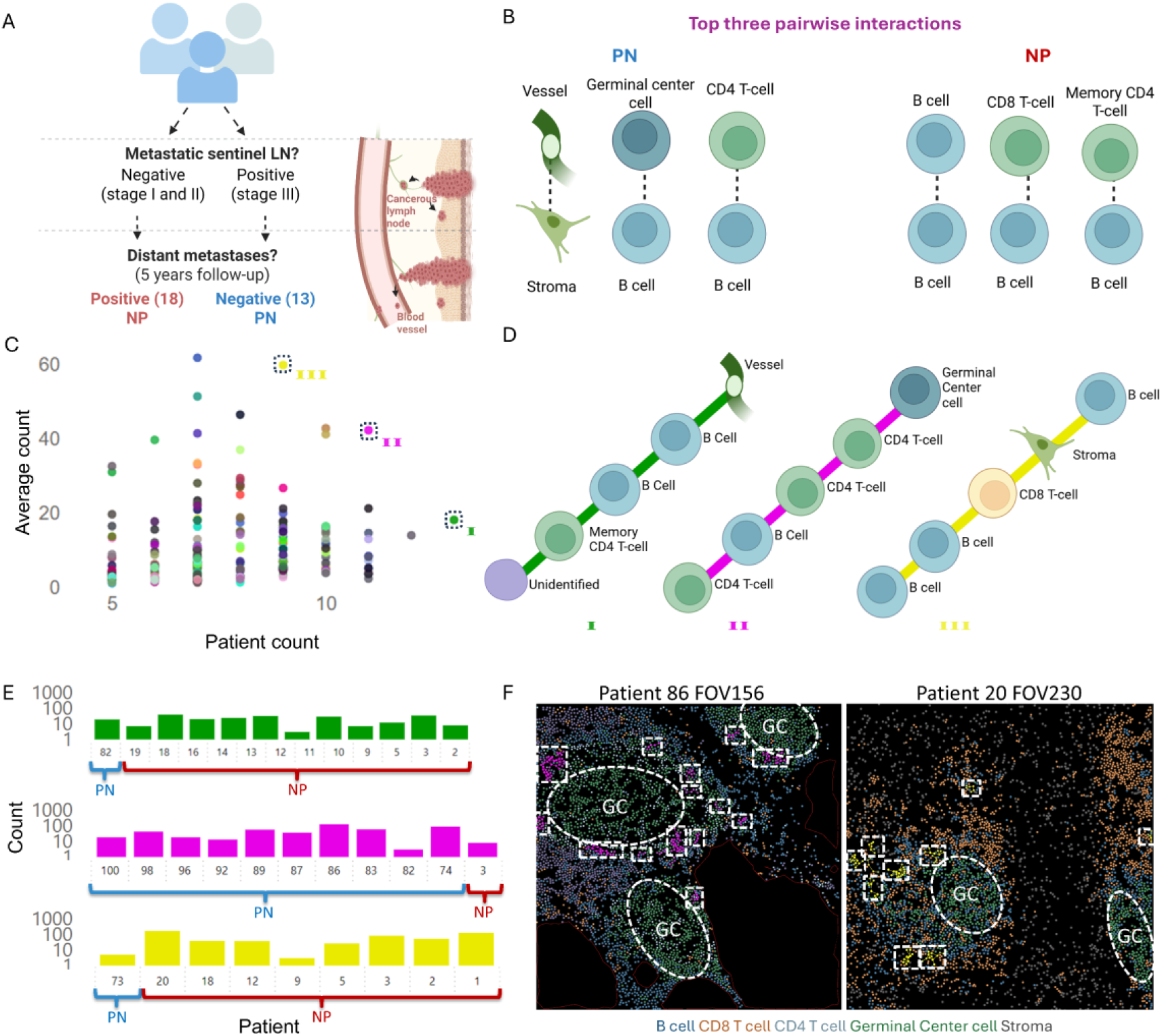
CISM analysis of NP versus PN. (**A**) The melanoma cohort includes 18 patients diagnosed with a primary tumor with no apparent tumor cells spreading to the lymph node (i.e., melanoma stage I-II) with widespread melanoma metastasis five years following treatment (Negative-Positive, NP) as described in Fig. 2A. The cohort was extended with 13 stage III melanoma patients who healed, i.e., patients with metastatic sentinel lymph nodes, and five years follow up prognosis of tumor-free lymph nodes and no metastases (Positive-Negative, PN). (**B**) The top three highly ranked motifs-induced pairwise interaction. The full distribution of motifs-induced pairwise interactions appears in Fig. S13A. (**C**) The mean per-patient number of discriminative motif instances (y-axis) in relation to the number of patients that contained that motif (x-axis). Each dot represents a discriminative motif (n = 210). Note motifs I (NP-associated, green), II (PN-associated, magenta), and III (NP-associated, yellow) that appeared in many patients and in many instances per patient. (**D**) The three discriminative motifs that were marked in C. The edge colors are consistent across the other figure panels. (**E**) The number of instances of motifs I-III (top-bottom, log scale) across patients. Only patients with at least one instance were listed. (**F**) Localizing motif II in the tissue of PN patient #86 (left) and motif III in the tissue of NP patient #20 (right). Each dot represents a cell colored according to its cell type: B cell, CD8 T-cell, CD4 T-cell, Germinal Center cells, and Stroma (other cell types are not shown). The germinal center (GC) is surrounded by a dashed white circle. Instances of motifs II and III are marked with dashed magenta and yellow rectangles (larger rectangles indicate multiple instances of the motif) respectively.

Focusing on changes in the organization of the lymph node immune microenvironment, we excluded all cells inside the tumor cell clusters in PN patients using a refined version of the convex hull algorithm (Kirkpatrick & Seidel, 1983, Methods). Cell type classification showed a similar cell type distribution for NP and PN patients (Fig. S12A). CISM analysis of NP versus PN found the optimal tradeoff between classification performance and sufficient discriminative motifs and their instances for interpretability in 5-cell motifs and discrimination stringency of 0.4 (Fig. S12B-C). CISM reached an AUC of 0.81, surpassing pairwise cell type machine learning with an AUC close to random (Fig. S12C – pink horizontal line) and GNN with an AUC of 0.76 (Fig. S12C – cyan horizontal line). The motif-induced pairwise interactions showed clear differences distinguishing between the disease states (Fig. 6B, Fig. S13). Most of the NP-associated discriminative motifs’ induced pairwise interactions appearing in either NP-vs-NN (Fig. 3C) or in NP-vs-PN (Fig. 6B), but not in both. The only motif-induced pairwise interaction that appeared in NP in both contexts was between B cells and CD8 T-cells, suggesting these interactions as putative specific markers for future metastases for stage II melanoma.

To interpret the NP-versus-PN discriminative motifs, we selected three prominent discriminative motifs that appeared in many patients with many instances per patient (Fig. 6C-D). Two of these motifs were NP-associated (motif I in green and III in yellow) and one was PN-associated (motif II in magenta) (Fig. 6E). Following the discriminative motifs’ localization at the boundaries of B cell zones that surrounded the germinal center in the context of NN-vs-NP (Fig. 5D), we examined the localization of NP-vs-PN discriminative motifs and identified two out of the three prominent motifs (II and III) that appeared near germinal centers (Fig. 6F). Motif II was PN-associated and appeared in 10/13 of PN patients adjacent to the germinal center, connecting it with the B cell follicles surrounding it (Fig. 6F). Motif III was NP-associated, included an edge between a B-cell and a CD8 T-cell and appeared in 6/18 of NP patients stereotypically at the edges of B cell follicles surrounding the germinal center (Fig. 6F, right). This latter specific B- and CD8 T-cell interaction and its spatial pattern aligned with our earlier observation in the context of NN-vs-NP (Fig. 5D). Altogether, interactions between B- and CD8 T-cells at the boundaries of B cell follicles surrounding the germinal center could be an early specific marker of future widespread metastasis in tumor-free lymph node stage II melanoma patients. The inference of a causal link which may lead to new targets is left to future studies.

### Generalizing CISM to reveal motifs associated with triple-negative breast cancer disease state

To demonstrate generalization, we applied CISM to analyze a MIBI-TOF-acquired human TNBC cohort consisting of 38 tissue sections of 37 patients (Keren et al., 2018). We partitioned the patients into short-term (less than 1000 days, N = 7 patients) and long-term (more than 1000 days, N = 30) survivors according to the time a person lived after diagnosis (Fig. S14A). Using the cell type classification (Fig. S14B) we applied CISM to extract 3- and 4-cell motifs and then selected those that discriminated between short- and long-term survivors with a smaller number of short-term associated discriminative motifs attributed to the imbalance between the disease states (Fig. S15). Unlike the melanoma cohort, in which the lymph nodes did not necessarily include tumor cells (NN and NP), in the TNBC cohort we did consider the tumor cells and their interactions with cells in the tumor microenvironment. Machine learning validation reached an AUC of 0.85, surpassing pairwise cell type machine learning and GNN (Fig. 7A, Fig. S16). Discriminative motifs-induced pairwise interactions revealed that long-term survivors were associated with the motif-induced pairwise interactions of DC/MONO cells with CD4 T cells, CD8 T cells with CD4 T cells, and between CD4 T cells, while the motif-induced pairwise interactions associated with short-term survival were of mesenchymal cells with CD8 T cells and Neutrophils with CD8 T cells (Fig. 7B, Fig. S17). These results propose interactions of CD4 T cells with other immune cells as a marker for long-term survival in contrast to interactions of CD8 T cells with neutrophils and mesenchymal cells. For spatial interpretation, we selected three discriminative motifs that were associated with long-term survivors (Fig. 7C-E), but did not find a clear stereotypic spatial localization of these motifs (Fig. 7F). Cumulatively, these results demonstrated that CISM is a general method to identify local cell structures associated with a disease state in single cell spatial data.

**Figure 7.**
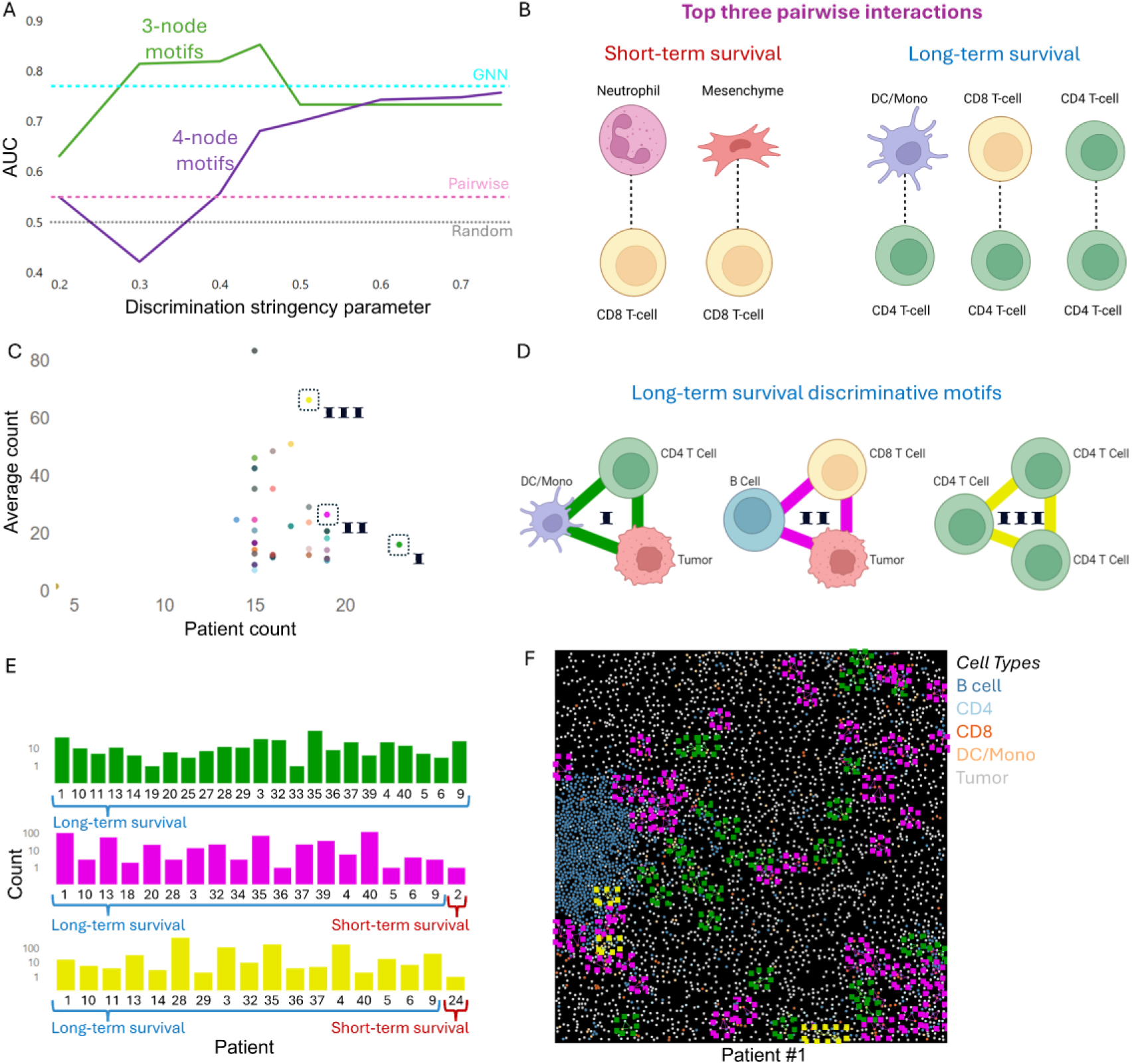
CISM analysis of triple-negative breast cancer cohort. (**A**) Machine learning validation of CISM’s discriminative motifs representation of TNBC short-term versus long-term survival discrimination. Leave one patient out cross-validation (LOOCV) model performance (AUC) (y-axis) over the discrimination stringency parameter (x-axis) (3-and 4-cell motifs in green and purple correspondingly). The pink, cyan, and black dashed horizontal lines represent the LOOCV AUC score of pairwise, GNN, and a random null model, correspondingly. The discriminative motifs derived from 3-cell motifs with the discrimination stringency parameter of 0.45 were used for further analysis. (**B**) The top three highly ranked interactions of motifs induced pairwise interaction (Fig. S17A). Note that short-term survival had a single discriminative motif and thus had only two motif-induced pairwise interactions. (**C**) The mean per-patient number of discriminative motif instances (y-axis) in relation to the number of patients that contained that motif (x-axis). Each dot represents a discriminative motif (n = 33). Note that motifs I (green), II (magenta), and III (yellow) are all associated with long-term survival and appeared in many patients and in many instances per patient. (**D**) The three discriminative motifs that were marked in C. The edge colors are consistent across the other figure panels. (**E**) The number of instances for motifs I-III (top-to-bottom, log scale) across patients. Only patients with at least one instance were listed. (**F**) Localizing motifs I-III in the tissue of the long-term survivor patient #1. Each dot represents a cell colored according to its cell type (right legend; other cell types are not colored). Instances of motifs I, II, and III are marked with dashed green, pink, and yellow rectangles correspondingly. Larger rectangles indicate multiple instances of the motif.

## Discussion

CISM is a two-step method for identification and characterization of fine-grained spatial inter-cellular modules associated with the physiological tissue states. First, using our FANMOD+ implementation, we can extract, in an unsupervised manner, a large set of spatial motifs in the patients’ multicellular spatial networks. Second, we select from the pool of motifs, in a supervised manner, the ones that are discriminative according to the question of interest. These two steps distill the discriminative motifs from the huge and noisy landscape of all putative sub-networks of intercellular interactions. These discriminative motifs can then be used for several downstream applications that we demonstrated in melanoma and TNBC: machine learning prediction of tissue’s physiological context, identifying “signature” discriminative motifs and motifs-derived cell distributions and pairwise interactions that associate with the physiological state, moving back to space to interpret the discriminative motifs’ stereotypic localization patterns in relation to coarser spatial scales. CISM outperforms cell type distributions, pairwise interactions and GNN representations in predicting disease tissue state. Moreover, CISM representation, that encodes the tissue state according to the fine resolution of a few cells and their local interactions, even surpassed an integrated multi-scale representation in predicting future melanoma metastases (Amitay et al., 2024), indicating that discriminative motifs can serve as powerful representations of tissue disease state. Intriguingly, we discovered that the exact spatial arrangement of the cells within the motifs, in terms of their specific cell-cell adjacencies (as indicators of potential interactions), contributed to the discrimination between different disease states. These results suggest that the local organization of a few cells in discriminative motifs are emergent properties that may define an intermediate spatial scale driving tissue function.

Although GNNs have shown great promise in classifying tissue states across multiple diseases (Tamir et al., 2024; Wu et al., 2022; Wu & Zou, 2022), their success comes at the cost of “black box” poor interpretability. This inherent difficulty in providing human meaningful explanations for the GNN’s decision hampers the possibility of generating specific mechanistic hypotheses. CISM approach of feature (i.e., discriminative motif) selection resembles the more interpretable classic machine learning approach, with “engineered” features that are easier to decipher. The exhaustive enumeration of all potential motifs followed by the selection of those motifs that discriminate between the disease states, provides these desired joint properties of discrimination power along with interpretability. Specifically, CISM offers several layers of interpretability (Fig. 1D). First, discriminative motifs-induced cell distribution and pairwise interactions highlight specific cell-cell interactions that, when considered in context, associate with the tissue state. Second, the ability to map discriminative motifs back to their location in the tissue space enables to link the two different spatial scales by characterizing the (fine scale) motif’s localization in relation to a (coarse scale) specific spatial context. We demonstrated these interpretability capacities by associating interactions between B and CD8 T-cells at the edges of B cell follicles surrounding the germinal center in melanoma NP patients when comparing to either PN or NN contexts. Some of the melanoma motifs included ”Unidentified” cells. While these cells did not have distinguishable markers in the protein data, concomitant transcriptomics could suggest that some of these may be plasmablasts (Amitay et al., 2024). The role of these interactions should be further evaluated in functional studies.

When applying CISM to TNBC, we associated long-term survival with interactions of CD4 T cells with dendritic cells aligning with a recent study suggesting that interactions between dendritic cells and T cells induce a more effective immune response (Shapir Itai et al., 2024). Moreover, the discriminative motif that appeared in the vast majority (23/30) of long-term survival patients was composed of edges between a CD4 T cell, a dendritic cell and a tumor cell, aligning with another study that argued that interactions between CD4 T cells and dendritic cells suppress tumor cells (Lei et al., 2023). For short-term survivors, we identified associated interactions between Neutrophil cells and CD8 T cells that were proposed as protumorigenic by inhibiting CD8 T cells proliferation (Duan et al., 2024) and more generally neutrophils have been previously suggested to modulate adaptive immune responses by inhibiting T cells (Aarts et al., 2019; Shafqat et al., 2023; Zhang et al., 2024).

Beyond enabling biological insight at the spatial scale of a few cells, CISM benefits from several additional advantages. The strategy of unsupervised motif enumeration followed by supervised context-dependent selection enables us to use the same motifs to answer different questions as shown for melanoma NP versus NN/PN. Moreover, this approach is suitable for characterizing differences in small cohorts, for example, in matched healthy-versus-disease tissue of the same single patient. However, CISM also has some limitations. The fine resolution of a few cells and their spatial arrangement inherently implies that CISM is less forgiving of errors in cell segmentation and in cell type classification. Also, CISM has multiple parameters that must be carefully configured. First, the cell type lineage resolution. Using more cell types provides better discrimination and more specific biological insight at the cost of combinatorially increasing the size of the motifs space. Thus, beyond increasing the required computational resources, this implies a dramatic decrease in the number and prevalence of discriminative motifs. Similar considerations take place regarding the choice of the motif’s size and the discrimination stringency parameter. Increasing these parameters should lead in principle to better discrimination, but is limited by the lower probabilities to generalize across patients. Practically, CISM users should balance these parameters according to quantitative readouts that include the number of discriminative motifs (Fig. 2C) and the discrimination performance (Fig. 2D) as a function of the discrimination stringency parameter. Automated parameter calibration is a possible direction for future research. The strict definition of a discriminative motif, of appearing in a sufficient fraction of patients of one disease state but excluded from all patients of the other disease state, could be too restrictive, especially with the expected increase in cohorts’ sizes in upcoming years, which inherently decreases the probability for the presence of discriminative motifs. This requirement for “hard” discrimination can lead to situations where motifs appear in many patients and even in many instances of patients of one disease state but also in very small fractions of patients and in small numbers of instances of the other disease state. These “almost discriminative” motifs can have great discriminative and interpretative value but are currently entirely excluded from our analysis. To remedy this limitation, CISM could incorporate a “softer” discriminative criterion such as the distance between distributions of motif frequencies of each disease state (e.g., the Wasserstein distance (Vaseršteĭn, 1969). The implementation of “soft“ discrimination could also be useful to extend CISM to multiple disease states and to incorporate motifs with finer resolution in their cell types and cell states. These ideas are left to future studies.

## Methods

### Experimental data

#### Melanoma MIBI-TOF

We analyzed an existing-acquired cohort of 51 human melanoma patients from Leeat Keren’s lab at the Weizmann Institute of Science (Amitay et al., 2024). The patients’ disease state was determined according to the clinical diagnostic of the sentinel lymph nodes (SLN) and then at least five years following treatment, except for one living patient who had a two year follow-up. Patients with or without the existence of tumor cells in their sLNs were defined as “positive” or “negative,” respectively. Patients who were diagnosed with the prognosis of widespread metastases in distant organs five years following treatment were defined as “negative-positive” (NP), while those who did not were defined as “negative-negative” (NN) or “positive-negative” (PN) according to their initial diagnosis. Cumulatively, we analyzed three patient groups: NN (N patients = 20, n tissue sections FOVs = 40), NP (N = 18, n = 36), and PN (N = 13, n = 42). The dataset includes 39-channel MIBI-TOF of sLNs with a resolution of 0.45X0.45 µm per pixel. In the original manuscript, each cell was classified into one of 26 cell types. We reduced the cell type resolution to ”tumor” and 15 non-tumor lymph node microenvironment cell types according to their lineage (Amitay et al., 2024) (Fig. S1) to increase the number of discriminative motifs and their occurrences. We included ”unidentified” cells in our machine learning analyses and excluded them from post-processing biological interpretability for selecting the three most abundant pairwise co-localized cell types (Fig. 3D, Fig. 6B) and for selecting an example for spatial arrangement analysis compared to the cell type (Fig. 4C).

#### TNBC MIBI-TOF

We analyzed a published MIBI-TOF-acquired human cohort TNBC dataset (Keren et al., 2018) that included 37 patients and 38 tissue sections with 38 protein channels and a spatial resolution of 0.5x0.5 µm per pixel. We used cell segmentation masks and assignment of each cell to one of 16 cell types as in the original study. The cell types used in this dataset include Tumor, Endothelial, Mesenchyme, Tregs, CD4 T cells, CD8 T cells, CD3 T cells, NK cells, B cells, Neutrophils, Macrophages, DC, DC/Mono, Mono/Neu, Immune other and Unidentified. We used the number of survival days since diagnosis as the clinical readout where patients who survived less than 1,000 days were defined as “short-term” survivors, and those who survived at least 1,000 days were defined as “long-term” survivors.

### Computational modeling

#### Defining the spatial multicellular network

We defined the spatial multicellular network, where single cells define the network nodes and are “colored” according to their cell type. Close-adjacent pairs of cells were connected with an edge using the Delaunay triangulation, excluding those edges between cells that distant more than 50 µm away from one another.

#### Excluding tumor regions from melanoma PN patients’ lymph node

In the melanoma cohort, we focused on disease-associated alterations of the lymph node immune microenvironment, specifically because NN and NP patients did not have tumor cells in their lymph nodes. Thus, when handling PN patients, we had to exclude tumor cells. We opted to exclude from the multicellular network all cells (tumor and non-tumor) in regions enriched with tumor cells to avoid potential tumor-immune interactions, as an obvious bias indicative of PN patients, which could serve as a “shortcut” for classification. Thus, we used the “alpha shape” (Kirkpatrick & Seidel, 1983), a generalization of the convex hull for a set of points in space (in our case - cell localizations) based on parameter α that controls the shape’s sensitivity. For α equal to zero, the alpha shape of a point set is the same as its convex hull. As α increases, the alpha shape shrinks with concave regions (derived from the point set) beginning to appear, gradually “cutting out” voids and indentations in the shape. We empirically set α to 0.01 after manual calibration (Fig. S18). Overall, we adjusted the multicellular network using the following pipeline: (1) identify all tumor cells, (2) divide them into connected components in the multicellular network, (3) calculate the alpha shape, and (4) exclude all cells inside the alpha shape from further analysis.

#### FANMOD+ implementation enables the extraction of large and multi-colored motifs

We adapt and extend the concept of “network motifs”, reoccurring sub-networks that define “building blocks” of complex networks (Milo et al., 2002), to characterize the fine-scale organization of multicellular spatial networks of human disease tissues. One of the important characteristics of the motif is a statistical measurement of its significance. A subgraph is considered a motif if its frequency in the observed graph precedes its frequency in random graphs generated by edge-switching that preserves the graph’s degree distribution. There are several existing tools to extract network motifs, but all have limitations regarding the motif size (i.e., number of cells) and the number of node “colors”, and graph type support (i.e., multigraph, directed/undirected graph). The only method that we found that supports node coloring is called FANMOD (Wernicke & Rasche, 2006). However, FANMOD has limitations on the number of colors as a function of the motifs’ size that it can analyze. Specifically, the maximum number of node colors that FANMOD can analyze is 15, and this number decreases as the motif size increases: at most, 15 colors for 4-node motifs and no coloring for 8-node motifs. This is a severe limitation for spatial single cell multiplexed data characterized by multiple cell types. We realized that this was an implementation (rather than algorithmic) limitation and thus adjusted the FANMOD source code, using the C++ boost library, to enable larger motif detection. We call our implementation FANMOD+ and it is publicly available at https://github.com/zaritskylab/FANMODPlus.

In more detail. To enumerate the set of all unique subgraphs within the network, Fanmod generates a unique representation of each subgraph, known as the “canonical representation”, that is invariant for different organizations of nodes and edges (i.e., isomorphic graphs are mapped to the same representation). Specifically, Fanmod is using the Nauty algorithm (McKay & Piperno, 2014) to generate a canonical representation that is encoded as a sequence of bits. FANMOD+ increases the space (i.e., number of bits) for the canonical representation from 64-bit fixed allocation to 128-bit dynamic allocation (i.e., physical memory allocation extended when used) (Schäling, 2011). Thus, FANMOD+ subgraph canonical representation enables the encoding of more colors and larger motifs, supporting the analysis of modern spatial multiplex single cell omics data. For instance, with 0-32 colors, FANMOD+ can support a maximal motif size of 5 nodes; with 128 colors, FANMOD+ can support a maximal motif size of 4 nodes. Note that increasing the motif size or the number of colors still has a computational cost (Fig. S19, description in the next section).

#### FADMOD+ benchmarking

To assess how the execution time increases as a function of more colors and larger motifs, we performed benchmarking using randomly generated graphs (Fig. S19). Since, on average for the Melanoma dataset, there were ∼10,000 cells in each patient’s sample, we generated random graphs with n = 10,000 cells, each represented as a pixel, randomly spread on a grid of size 10,000 x 10,000 pixels and connected nearest neighbors cells using Delaunay triangulation with ∼30,000 edges (∼3n following the Euler rule for planar graph when n ≥ 3). We generated five random graphs for each combination of motif size (3-5) and colors (in the range of 8-128). Due to RAM usage limitations of 64GB, we limited our benchmarking to five nodes and 32 colors. The benchmarking was executed on an AMD Ryzen 9 5900HX CPU with 64GB RAM. The average runtimes for 3 and 4 -node motifs with 128 colors were 30 seconds and 1846 seconds (i.e., ∼30 minutes) correspondingly. For 5-node motifs with 32 colors, the average runtime was 1539 seconds (full results Fig. S19). Overall, the runtime increases exponentially with the number of colors.

#### Extracting network motifs with FANMOD+

We applied FANMOD+ to each patient’s tissue sample to extract and enumerate thousands, tens-of-thousands and hundreds-of-thousands of motifs of size 3-to-5 cells from the space of hundreds-of-thousands, a million and a few millions candidate subgraphs, correspondingly (Table S1). We calculated each n-cells motif’s frequency with respect to the total number of corresponding subgraphs of size *n* in the patient’s sample. We used 1000 random graphs for the statistical test and included only the significant motifs for the analysis (p-value < 0.05). To illustrate the definition of what defines a subgraph as a motif we provide a specific example. Consider a network in a single field of view that contains 10,000 nodes (cells), each with 1 of 10 colors (cell types) and 30,000 edges (cell-cell interactions) (euler rule for planner graph). For a 4-node subgraph with 3 edges each, the probability for a given cell type composition is 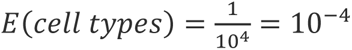, and the probability for the cells arrangement is a subgraph is 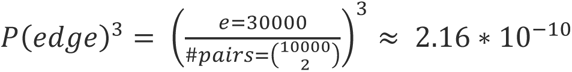. Thus, the estimated expected number of subgraph instances is 𝐸(4 𝑛𝑜𝑑𝑒 𝑐𝑎𝑛𝑑𝑖𝑑𝑎𝑡𝑒𝑠) ∗ 𝑃(𝑠𝑡𝑟𝑢𝑐𝑡𝑢𝑟𝑒) ∗ 𝑃(𝑐𝑒𝑙𝑙 𝑡𝑦𝑝𝑒𝑠) ≈ (_4_^10000^) ∗ 2.16 ∗ 10^−10^ ∗ 10^−4^ ≈ 4.17 ∗ 10^14^ ∗ 2.16 ∗ 10^−10^ ∗ 10^−4^ ≈ 9. Thus, in this specific example, the number of instances in the original network would need to significantly exceed 9 instances to be considered a motif.

#### Selecting discriminative motifs

*Context-dependent discriminative motifs* are those motifs that sufficiently appear in patients of one physiological context and do not appear at all in patients of the other physiological context. The degree of presence of a motif in each context is measured as the fraction of patients in that context that had at least one instance of that motif. Discriminative motifs were selected based on a tunable *discrimination stringency parameter* that defines a threshold on the degree of presence in a given context with the constraint that the motif does not appear in any of the patients of the other context.

#### Ranking discriminative motifs

To reduce the computational complexity, improve interpretability and remove redundant features, we selected the top 𝑛_𝑑𝑖𝑠𝑐_𝑚𝑜𝑡𝑖𝑓𝑠_ discriminative motifs in each leave-one-out iteration. The ranking of the discriminative motifs was determined according to their frequencies - the ratio between the number of motif instances to the total number of corresponding subgraphs (of the motif’s size) in the patient’s sample. Our assumption was that motifs appearing in more patients and in more instances per patient are a preferable representation of the physiological context.

#### Machine learning evaluation of the discriminative motifs representation

To assess the discriminative capacity of motifs’ representation of the disease tissue state, we performed Random Forest-based leave-one-patient-out cross-validation for binary classification between two disease contexts. The reason for using a Random Forest model was its simplicity and stable performance even with default settings. The reason for the leave-one-patient-out cross-validation was the small cohorts sizes that are usually limited to tens of patients. Thus, we performed iterations of training and testing, each iteration with one patient at the test and the rest used for training. Each iteration followed the processes depicted in Fig. 1C (left-to-right): (1) keeping one patient out, (2) extracting motifs from all remaining patients, (3) selecting discriminative motifs and ranking the top *n* discriminative motifs (i.e., 𝑛_𝑑𝑖𝑠𝑐_𝑚𝑜𝑡𝑖𝑓𝑠_), (4) representing a patient by the (sparse) feature vector encoding the per-patient frequencies of these highest ranked discriminative motifs. We used the Random Forest binary classification model with default parameters because of its simplicity, efficient and fast execution, and being suitable for interpretability. From the aggregated patient classification probabilities, we computed the receiver operating characteristic curve (ROC) and calculated the corresponding area under the curve (AUC) score. We set the number of features (𝑛_𝑑𝑖𝑠𝑐_𝑚𝑜𝑡𝑖𝑓𝑠_) to *n = 30* in all classification tasks throughout this manuscript because it provided a nice balance between stable performance and rich representation. By stable performance, we refer to a representation that was not susceptible to fluctuations in the AUC scores as a function of the discrimination stringency parameter that typically occurs when using a small number of features. By rich representation, we refer to a sufficient number of discriminative motifs for high discriminative performance along with a manageable number of discriminative motifs for downstream evaluation and interpretation. Since the heuristic approach of ranking the motif’s importance and selecting a subset to train the model, the AUC as a function of the discrimination stringency parameter is not necessarily monotonic.

#### Permutation test of the machine learning performance

To assess if the prediction results and the selected features are significant and not overfitted to the data, we conduct a permutation test by permutating the patient’s disease states, i.e., their classification labels. We executed the complete evaluation pipeline and measured the ROC-AUC score for each permutation. We did not consider permutations that led to no discriminative motifs in at least one of the iterations. This permutation statistical test’s p-value was determined as the ratio between the number of permutations that resulted with AUC scores higher than the observed labels, and the total number of permutation trials.

#### Machine learning evaluation of cell type distribution

We used machine learning to evaluate the discriminative performance of cell type distribution. We followed the same process, i.e., the Random Forest leave-one-out-cross-validation. We computed the receiver operating characteristic curve (ROC) from the aggregated patient classification probabilities and calculated the corresponding area under the curve (AUC) score.

#### Explicit pairwise interactions

We measure the probability of an edge connecting two cell types across patients for each disease state. First, we created a pairwise cell-type interactions matrix with the probability of an edge connecting to two cell types for each field of view (FOV) from a patient at the given disease state:

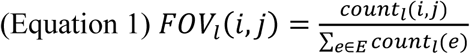

Where 𝑙 is a field of view, E is the set of all the cell-type interactions, L(p) is the patient’s FOVs and 𝑖, 𝑗 are cell types. Then, we summed the overall frequencies across the field of views and normalize:

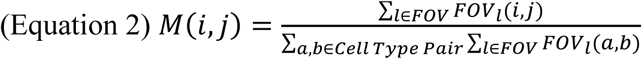

Where 𝑙 is a field of view and 𝑎, 𝑏, 𝑖, 𝑗 are cell types.

#### Machine learning evaluation of pairwise cell-cell interactions

We used machine learning to evaluate the discriminative capacity of pairwise interactions. We represented a patient with the probabilities of the pairwise cell-type interactions:

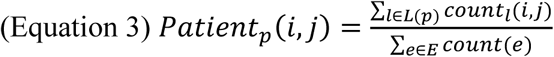

Where 𝑙 is a field of view, L(p) is the patient’s FOVs and 𝑖, 𝑗 are cell types. We followed the Random Forest leave-one-out-cross-validation. From the aggregated patient classification probabilities, we computed the receiver operating characteristic curve (ROC) and calculated the corresponding area under the curve (AUC) score.

#### Discriminative motifs-induced cell type distribution

The discriminative motif-based cell type distribution was derived from the pool of discriminative motifs. For a given disease state we pooled all the discriminative motif instances across patients’ field of views (FOVs) from that disease state and calculated the motifs-induced cell type distribution:

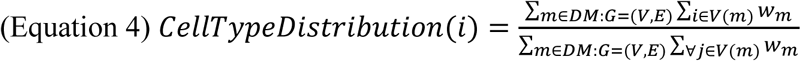

Where *i* is cell type, *m* is a discriminative motif, *DM* is the set of all discriminative motifs derived from all patients of a given disease state, and 𝑤_𝑚_ is the total count of discriminative motif instances across patients.

#### Discriminative motifs-induced pairwise interactions

The discriminative motif-based pairwise interactions were derived from the pool of discriminative motifs. For a given disease state we pooled all the discriminative motif instances across patients’ field of views (FOVs) from that disease state and calculated the motifs-induced pairwise interactions:

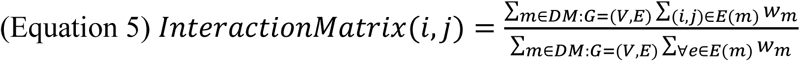

Where *i* and *j* are cell types, *m* is a discriminative motif, *DM* is the set of all discriminative motifs derived from all patients of a given disease state, and 𝑤_𝑚_ is the mean motif frequency across patients.

### Measuring the contribution of the motif’s spatial arrangement to disease state prediction

To measure the contribution of the motif structure compared to only considering the cell type composition, we replaced each discriminative motif with its corresponding cell type composition vector and used machine learning to evaluate the derived cell type composition representation in comparison to the corresponding motifs’ representation. Next, we incorporated motif information back to the cell type composition representation, one motif at a time, to evaluate the residual contribution of each motif to the cell type composition. More specifically, the cell type composition vector derived from a motif was represented as a vector of a fixed length according to the number of cell types, where each entry holds the number of instances of the corresponding cell type in the motif. For example, cell type composition I in Fig. 4C (green column) is composed of a B-cell, a HEVs cell, and two Memory CD4 T-cells. Note that different motifs can be mapped to the same cell type composition. For instance, two motifs with different inner structures that comprise 2 * x, 1 * y, and 2 * z cell types are mapped to the same cell type composition vector of (2, 1, 0, 2), given cell types (x, y, t, z). Thus, the evaluation was performed as follows. First, we replaced all discriminative motifs with their corresponding cell type composition vectors and measured the AUC of the cell type composition representation using Random Forest leave-one-patient-out as the baseline. Note that each cell type composition can correspond to multiple motifs, where some of these motifs do not meet the strict discrimination criterion used to define the discriminative motifs.

Second, starting from this cell type composition representation, we introduced back the spatial information of one cell type composition by replacing the cell type composition feature with its corresponding discriminative motifs. This replacement induced a hybrid representation that contains both cell type composition and discriminative motif features. This representation was used for machine learning assessment, where the classification gain with respect to the cell type composition baseline was attributed to the contribution of the corresponding motifs’ spatial arrangement. This process was iteratively repeated for each cell type composition, that were then ranked according to the contribution of the corresponding motifs’ spatial arrangement. The difference between the residual gain and the baseline score captures the contribution of the specific set of discriminative motifs for prediction compared to the generalization of the cell type composition.

### Machine learning evaluation of the discriminative cell type composition representation

To evaluate the motifs cell type composition information without the specific inner structure, we repeated the steps of context-dependent motifs’ selection and machine learning analysis, but instead of using the discriminative motifs, we used the discriminative cell type composition representations, i.e., cell type compositions that sufficiently appear in patients of one physiological context and do not appear at all in patients of the other physiological context. Each patient was represented by the (sparse) vector of its cell type composition frequencies. For instance, if multiple motifs are mapped to a single cell type composition, then the frequency is calculated as overall number of instances across the corresponding motifs count with respect to the total corresponding subgraphs. The cell type composition representation is less specific than the motif representation and, therefore, expects to generate a smaller number of discriminate cell type compositions.

### Graph neural network (GNN) disease state prediction

We performed a whole graph classification to predict the disease state using a two-layer graph convolutional network (GCN) (Kipf & Welling, 2016), and a fully-connected layer at the end (classification head). The spatial multicellular network defines the input for the GCN where the attribute of each node was its corresponding cell type. We aggregated the information across nodes after the final classification layer, using global pooling (average between the global max pooling and the global mean pooling), to get a single disease state prediction per sample. The model was optimized with the Adam optimizer (Kingma & Ba, 2014) and using the cross-entropy loss function:

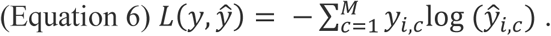

We performed leave-one-patient-out cross-validation. The training was as follows: each patient was held out for test, and the remaining samples were split into train-validation sets (75% train, 25% validation). After 500 epochs, we selected the optimal weights according to the epochs’ validation losses, and we used the corresponding model for the test. The reason for the train-validation-test splits was to avoid overfitting due to the relatively small dataset.

### Code and data availability

The CISM Python source code is publicly available at https://github.com/zaritskylab/CISM, and the FANMOD+ C++ source code is publicly available at https://github.com/zaritskylab/FANMODPlus. The melanoma dataset (V1) will become publicly available before journal publications. The TNBC processed dataset is publicly available at https://github.com/zaritskylab/CISM/blob/main/analysis/Tutorial/mibitof_tnbc_pred_cell_type.csv (Keren et al., 2018).

## Funding and Acknowledgments

This research was supported by the Wellcome Leap ΔTissue program (to AZ). L.K. holds the Fred and Andrea Fallek President’s Development Chair and is supported by the Enoch foundation research fund, the Abisch-Frenkel foundation, the Rising Tide foundation, the Sharon Levine Foundation, Fundación Alberto Palatchi, Dwek center for cancer immunotherapy, and grants from the European Research Council (948811), Israel Science Foundation (2481/20), the Rosetrees Foundation (10004), the Israel Precision Medicine Partnership Program (3830/21), and the Melanoma Research Alliance Team Science Award (1200724).

## Author Contribution

A Zaritsky and A Zamir conceived the study. A Zamir developed the computational method and analyzed the data. YT performed the GNN analysis and developed visualization tools. YA and LK generated and preprocessed the data. A Zamir and A Zaritsky interpreted the data and drafted the manuscript, with the help of YA and LK. All authors edited the manuscript and approved its content. LK mentored YA. A Zaritsky mentored A Zamir and YT.

