## Supplementary Information for "Context-dependent spatial multicellular network motifs for single-cell spatial biology"

### Supplementary Figures

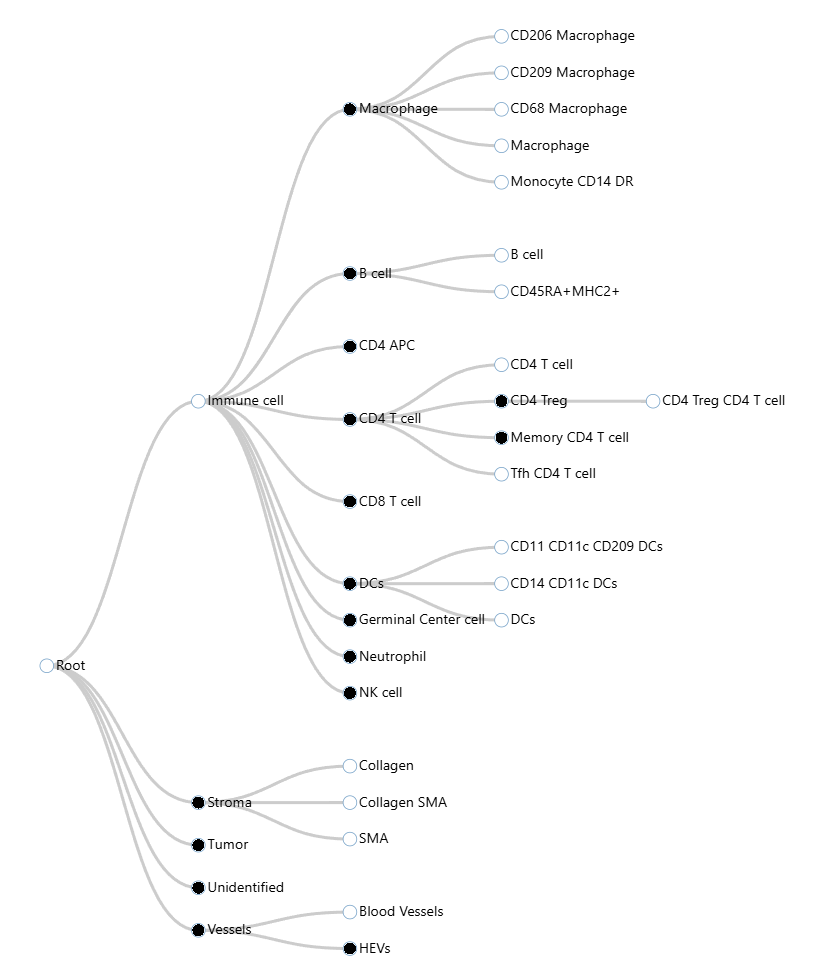

**Figure S1. Melanoma cell type hierarchy.** The hierarchy tree’s leaves (n = 26) represent the full-resolution cell types used in (Amitay et al., 2024). Black filled nodes (n = 16) represent the cell types used in this study as input to CISM.

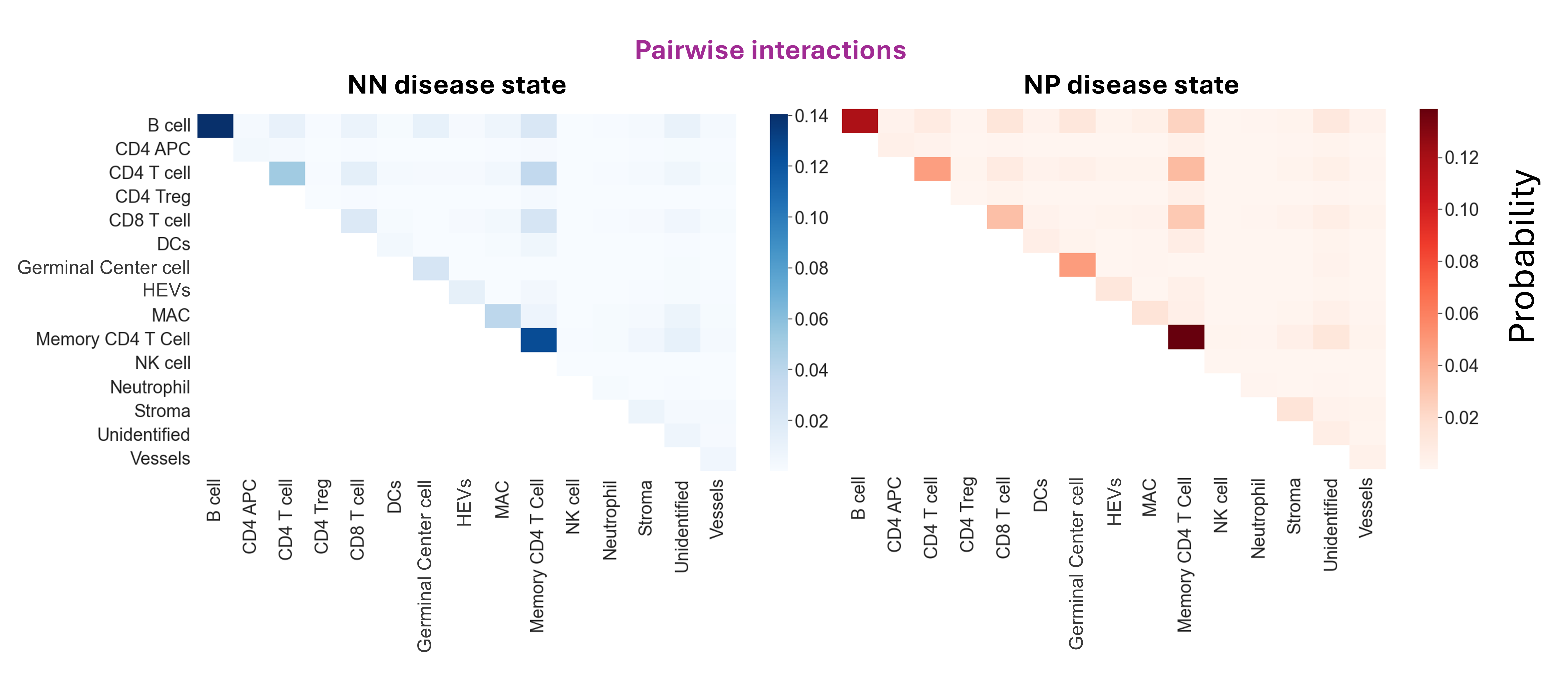

**Figure S2. The distribution of pairwise interactions**. The cell type pairwise edge probability in the cohorts’ patients pooled set of multicellular networks (without using motifs). Spearman correlation between NN and NP was 0.95 with p-value ≤ 0.0001.

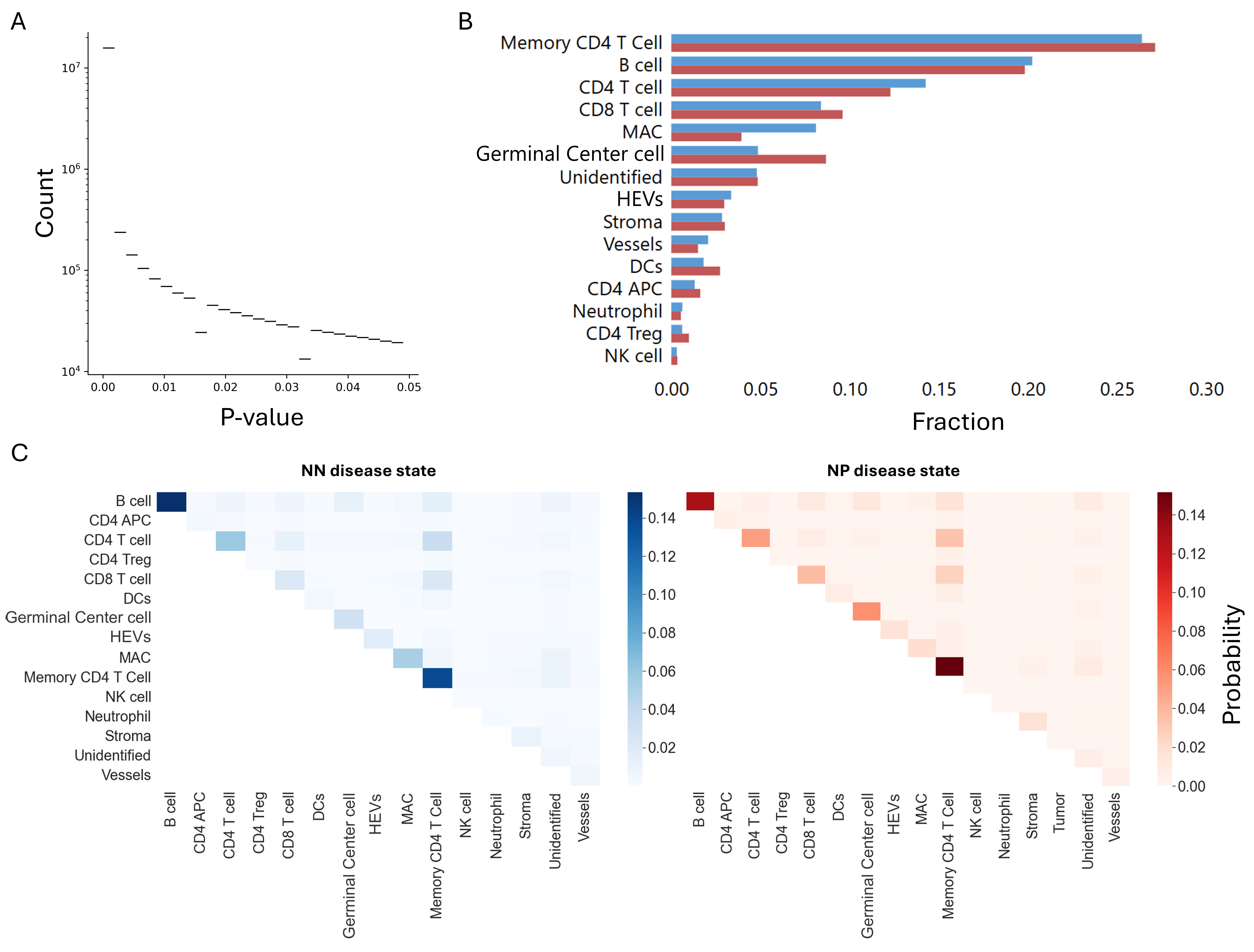

**Figure S3. Motifs-induced cell type and pairwise distributions in NN and NP disease states.** (**A**) The number of different 3-4-5 motifs (y-axis, log scale) for p-value intervals (x-axis). (**B**) Motifs-induced distribution of cell types pooled across all 4-cell motifs from all NN and NP patients. (**C**) Motifs-induced distribution of pairwise interactions. The cell type pairwise edge probability in the cohorts’ patients pooled set of 4-node motifs.

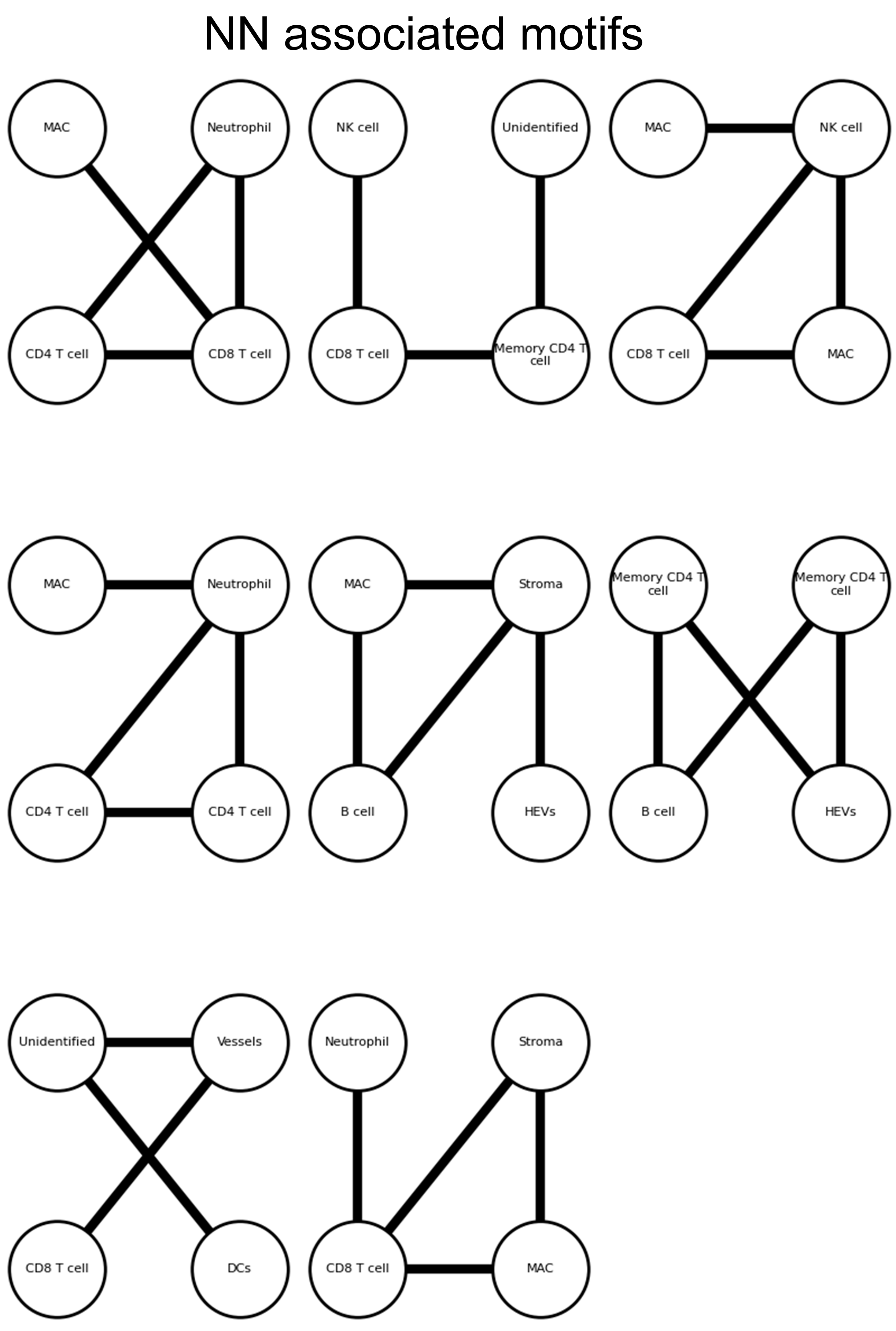

**Figure S4.** The eight 4-node NN-associated discriminative motifs in the context of NN versus NP that were used for the classification in Fig. 2D with the selected discrimination stringency parameter.

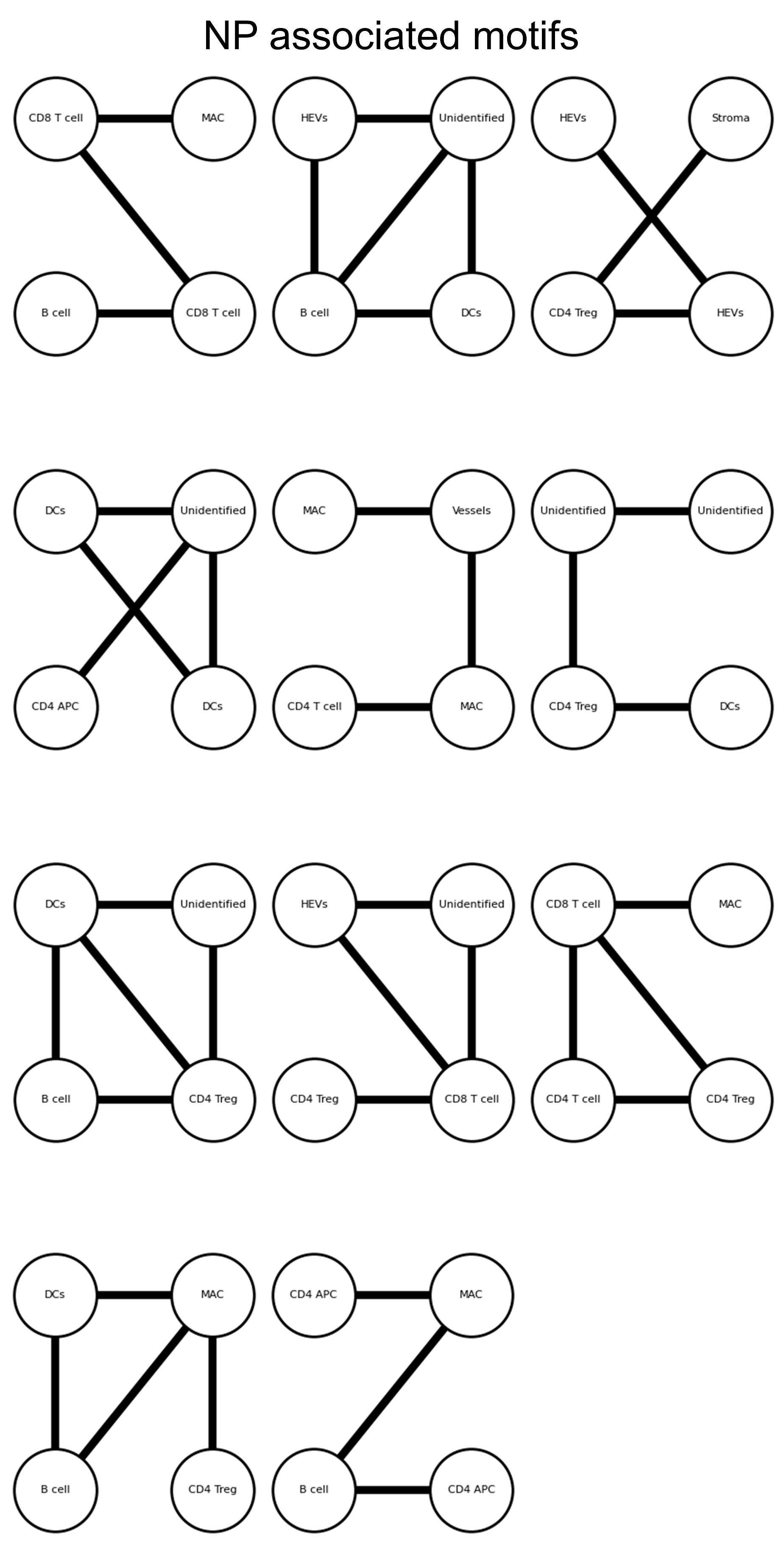

**Figure S5.** The eleven 4-node NP-associated discriminative motifs in the context of NN versus NP that were used for the classification in Fig. 2D with the selected discrimination stringency parameter.

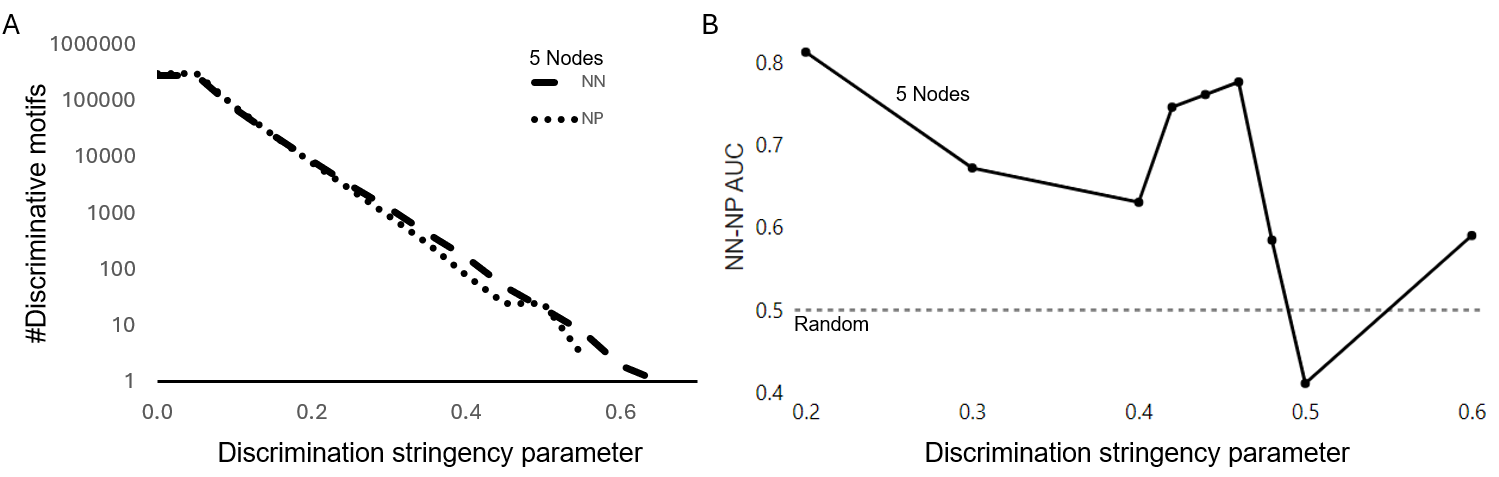

**Figure S6. CISM analysis of NN versus NP with 5-node motifs.** (**A**) The number of 5-cell discriminative motifs (y-axis, log scale) over the discrimination stringency parameter (x-axis). (**B**) Machine learning validation of melanoma disease state discrimination with CISM’s discriminative motifs representation. Leave one out cross-validation (LOOCV) model performance (AUC) (y-axis) over the discrimination stringency parameter (x-axis). The dropped AUC for stringent discrimination criteria is caused by training overfitting due to the small number of discriminative motifs per leave-one-out iteration.

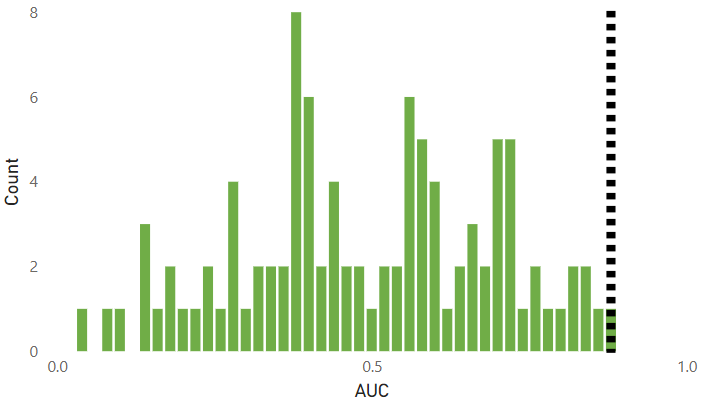

**Figure S7. Patient disease state permutation test in the context of NN versus NP.** AUC histogram of 100 random permutations of the patients’ disease state. Three permutations were excluded from the plot because they did not include discriminative features. The dashed black line is the control (unpermuted) AUC result of 0.88 leading to a p-value = 0.01.

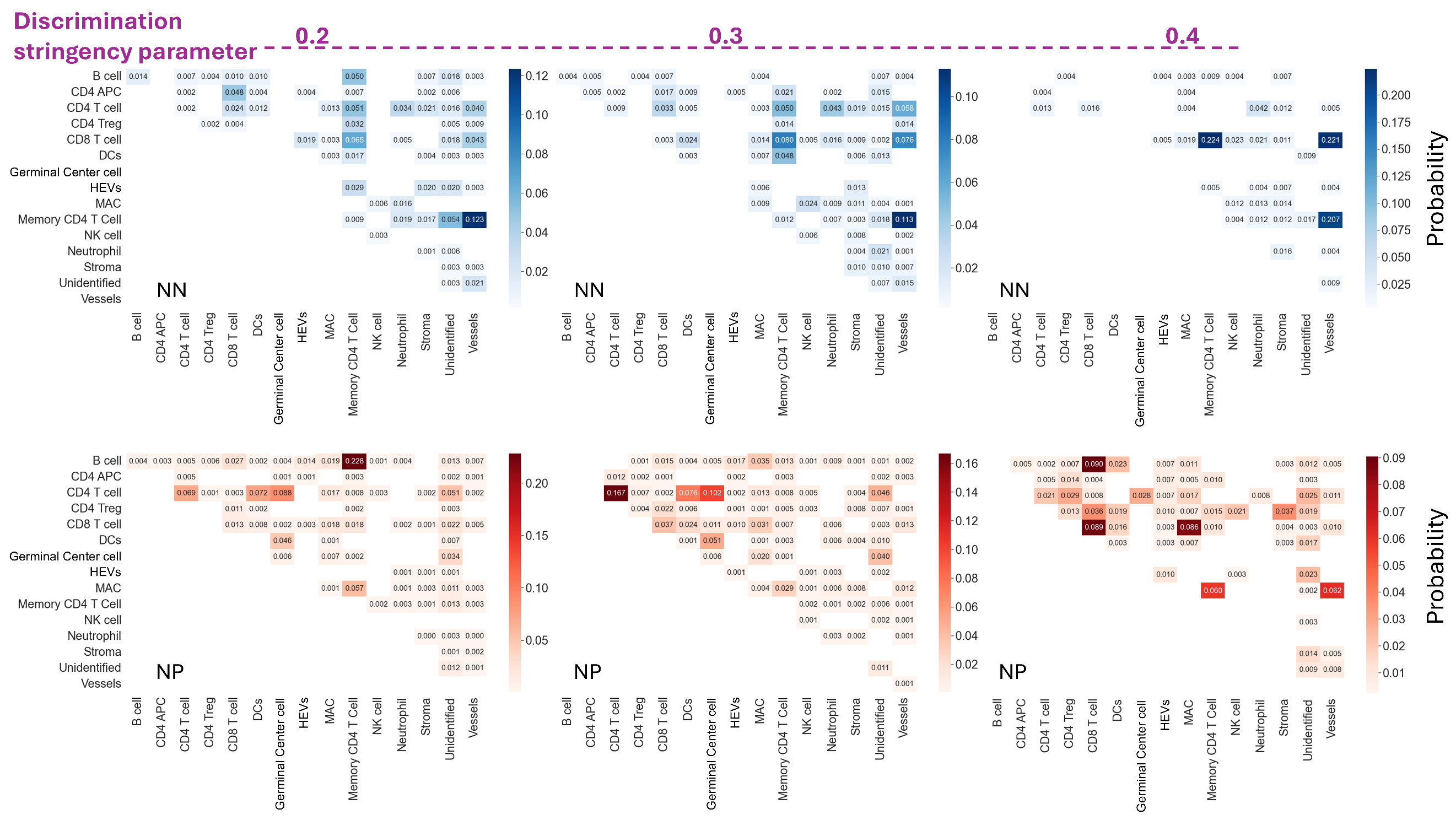

**Figure S8. Robustness of the discriminative motifs-induced pairwise interactions to the discrimination stringency parameter.** Cell type pairwise edge probabilities in the pooled set of instances of 4-cell discriminative motifs associated with NN (1^st^ row) and NP (2^nd^ row) across all patients for discrimination stringency parameter values of 0.2, 0.3, or 0.4 (left-to-right). The number of discriminative motifs in NN and NP was 56 and 100 correspondingly for discrimination stringency parameter of 0.2, 49 and 84 correspondingly for 0.3, and 16 and 34 correspondingly for 0.4. The Spearman correlation between the pairwise interaction matrices across consecutive thresholds was higher compared to the correlation attained with the same threshold across disease states. The correlation between consecutive thresholds: corr(NN[0.2], NN[0.3]) = 0.6248, p-value ≤ 0.0001; corr (NN[0.3], NN[0.4]) = 0.4160, p-value ≤ 0.0001; corr(NP[0.2], NP[0.3]) = 0.6833, p-value ≤ 0.0001; corr(NP[0.3], NP[0.4] = 0.3599, p-value ≤ 0.00017); The correlation between disease states using the same threshold: corr(NN[0.2], NP[0.2]) = 0.2785, p-value ≤ 0.0041; corr(NN[0.3], NP[0.3]): corr = 0.0424, p-value ≤ 0.6672; corr(NN[0.4], NP[0.4]) = 0.0682, p-value ≤ 0.4894.

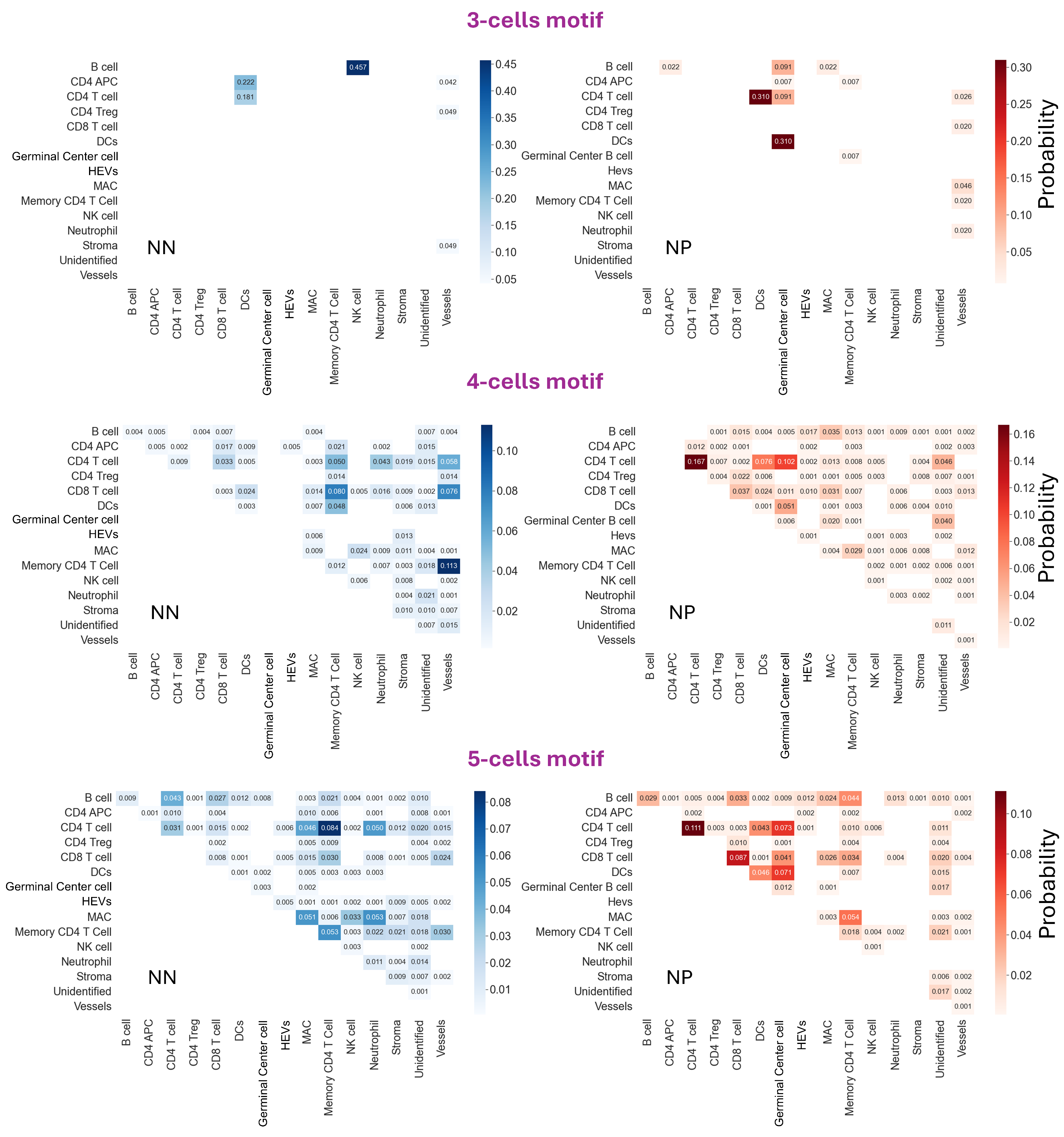

**Figure S9. Stability of discriminative motif-induced pairwise interactions to the motif size.** Cell type pairwise edge probabilities in the pooled set of instances of discriminative motifs associated with NN (1^st^ column) and NP (2^nd^ column) across all patients for 3-, 4-, 5- cell motifs. The discrimination stringency parameter was set to 0.3, which was the most stringent discrimination parameter that did not induce zero discriminative motifs. The number of discriminative motifs in NN and NP was 4 and 7 correspondingly for 3-cell motifs, 49 and 84 correspondingly for 4-cell motifs, 137 and 102 correspondingly for 5-cell motifs. The Spearman correlation between the pairwise interaction matrices across consecutive motif sizes was higher compared to the correlation attained with the same motif size across disease state. The correlation between consecutive motif sizes: corr(NN[3-cells], NN[4-cells]) = 0.0530, p-value = 0.5910; corr(NN[4-cells], NN[5-cells]) = 0.4872, p-value ≤ 0.0001; corr(NP[3-cells], NP[4-cells]) = 0.1624, p-value ≤ 0.0978; corr(NP[4-cells], NP[5-cells]) = 0.6145, p-value ≤ 0.0001; The correlation between disease state with the same motif sizes: corr(NN[3-cells], NP[3-cells]) = 0.0390, p-value ≤ 0.6924; corr(NN[4-cells], NP[4-cells]) = 0.0424, p-value ≤ 0.6673; corr(NN[5-cells], NP[5-cells]) = 0.2582, p-value ≤ 0.0079. Note that the poor correlation between 3- and 4-cell motifs was due to low number of discriminative motifs with 3-cell motifs (Fig. 2C).

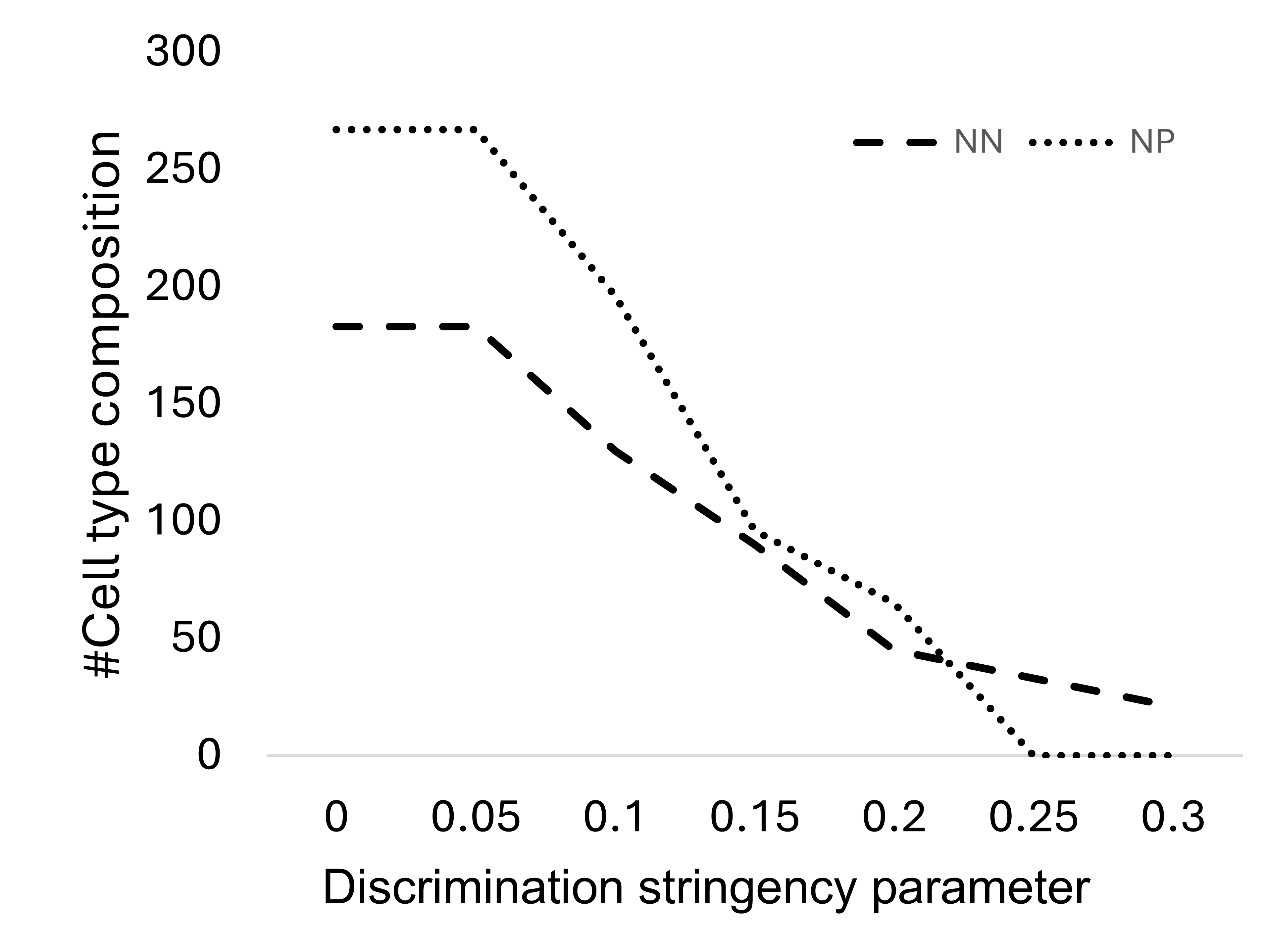

**Figure S10. A number of discriminative cell type composition representation.** The number of 4-cell discriminative cell type composition (y-axis) over the discrimination stringency parameter (x-axis).

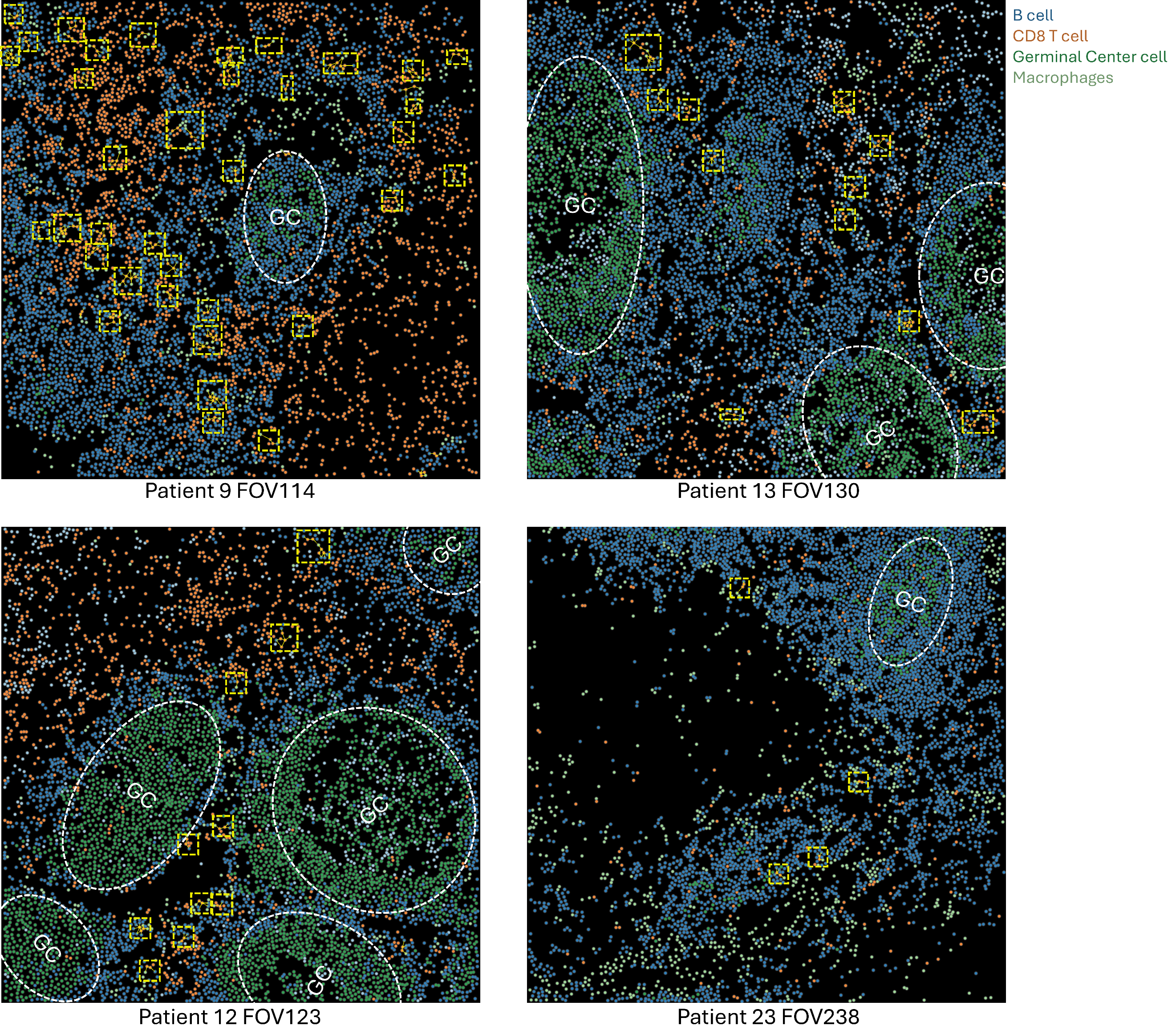

**Figure S11. Stereotypical localization of NP-associated motif III instances at the edges of B cell follicles that surround the germinal center.** NP patients #9 (top left), #13 (top right), #12 (bottom left), and #23 (bottom right). Each dot represents a cell colored according to its cell type: B cell, CD8 T cell, Germinal Center cells, and macrophages (other cell types are not shown). The germinal center (GC) is surrounded by a dashed white circle. Instances of motif III (Fig, 5B) are marked with dashed yellow rectangles (larger rectangles indicate multiple instances of the motif). The localization pattern aligns with Fig. 5D.

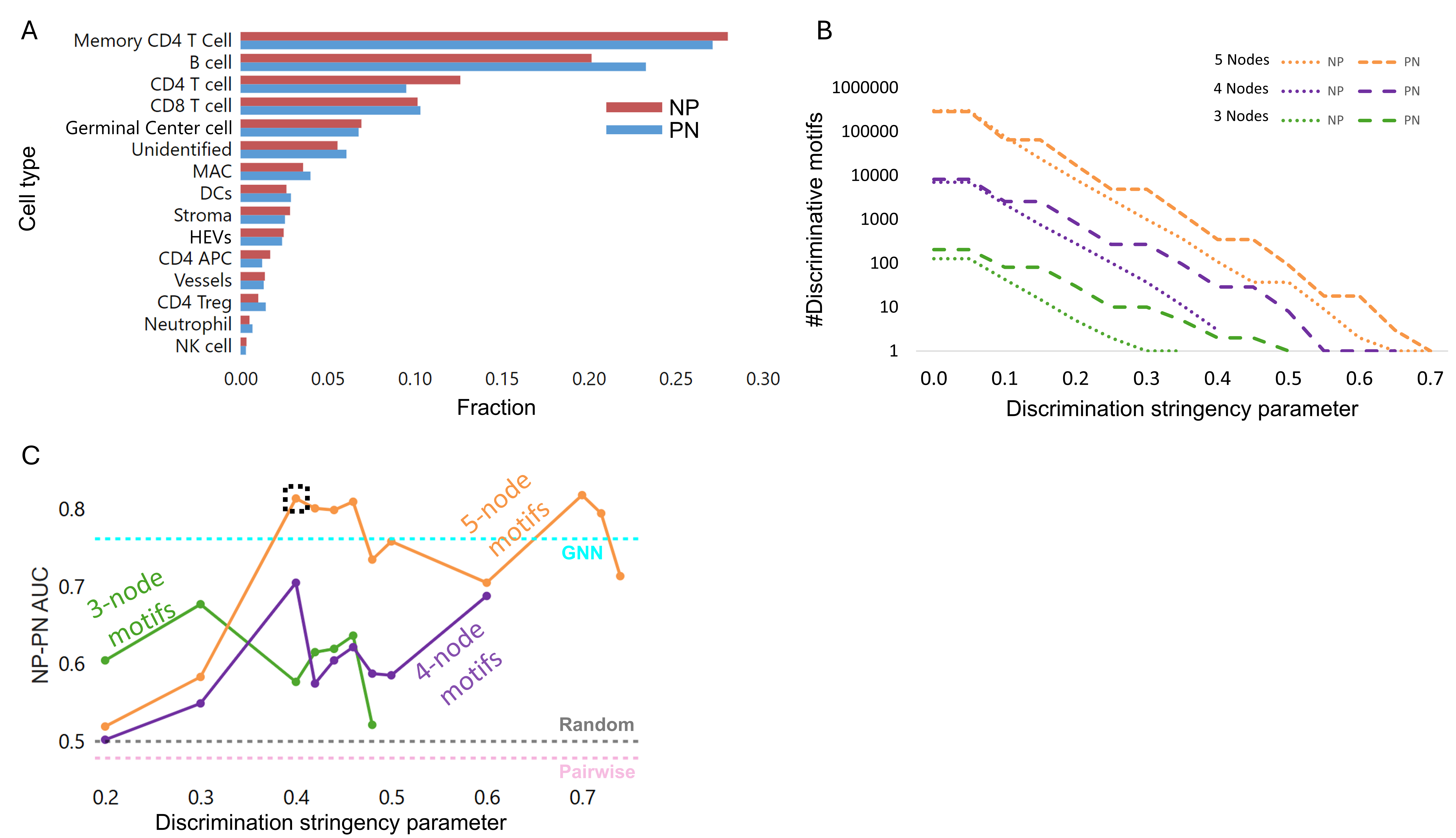

**Figure S12. CISM classification of PN-versus-NP patients.** (**A**) Cell type distribution for PN and NP patients. (**B**) The number of 3-cell (green), 4-cell (purple), and 5-cell (orange) context-dependent discriminative motifs (y-axis, log scale) over the discrimination stringency parameter (x-axis). (**C**) Machine learning validation of melanoma disease state discrimination with CISM’s discriminative motifs representation. NP-versus-PN leave one patient out cross-validation (LOOCV AUC, y-axis) over the discrimination stringency parameter (x-axis) (green - 3-cell motifs, purple- 4-cell motifs, orange- 5-cell motifs). The pink, cyan, and black dashed horizontal lines represent the LOOCV AUC score of pairwise, GNN, and a random null model, correspondingly. The discriminative motifs derived from 5-cell motifs with the discrimination stringency parameter of 0.4 for discriminative motifs selection, that best balances discrimination, generalization and interpretability, were used for further analysis.

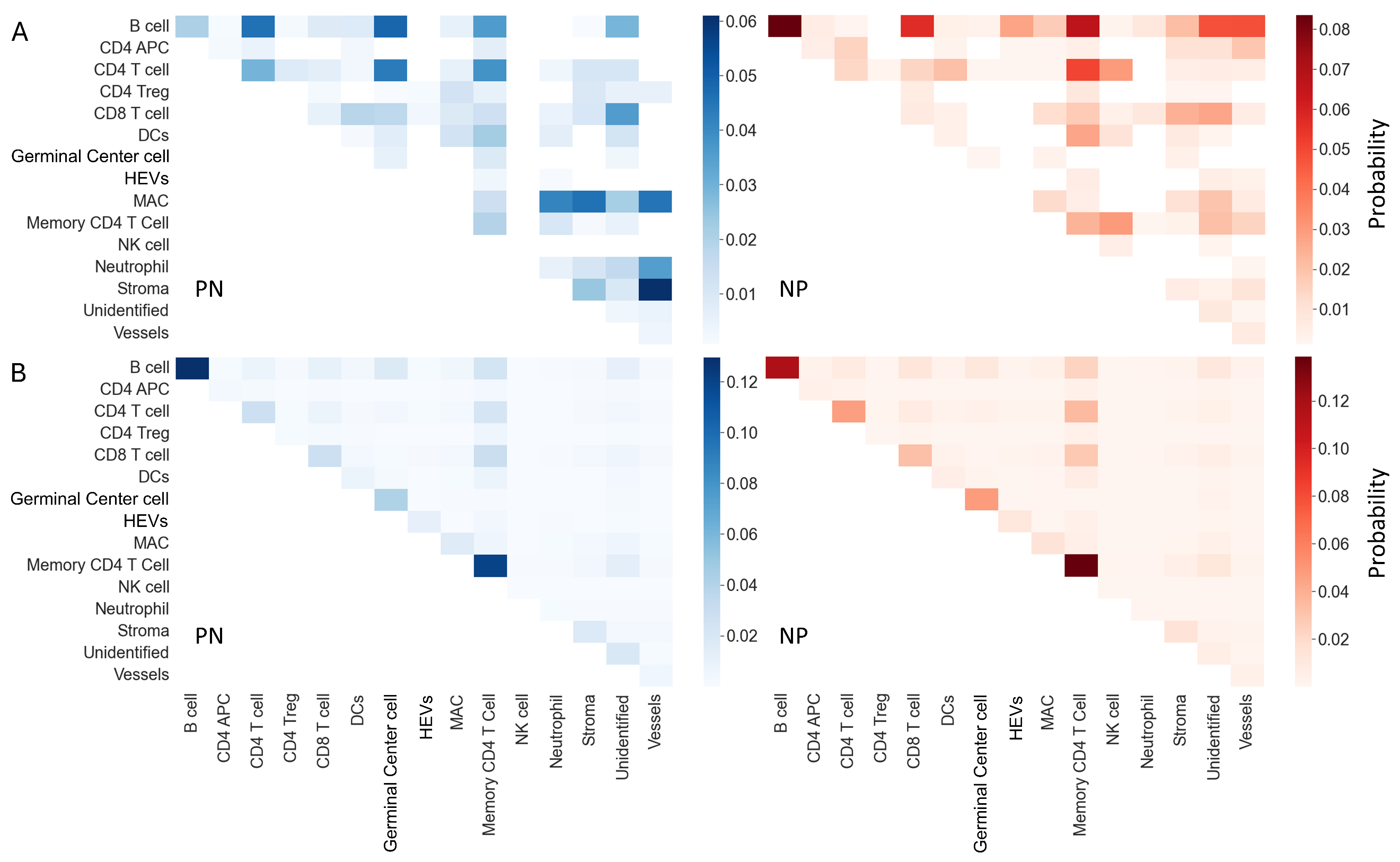

**Figure S13. Motifs-induced pairwise interactions in the context of PN versus NP.** (**A**) The distribution of motifs-induced pairwise interactions. The probability of two cell types to be connected with an edge in the cohorts’ patients pooled set of instances of discriminative motifs associated with NN patients (left**,** n = 120 motifs) and NP patients (right**,** n = 90 motifs). (**B**) Explicitly enumerating the distribution of pairwise interaction for PN versus NP patients according to the probability of two cell types being connected with an edge in the multicellular network did not show a clear distinction in the context of disease state. The distribution of pairwise interactions. The probability of two cell types to be connected with an edge in the cohorts’ patients pooled set of multicellular networks.

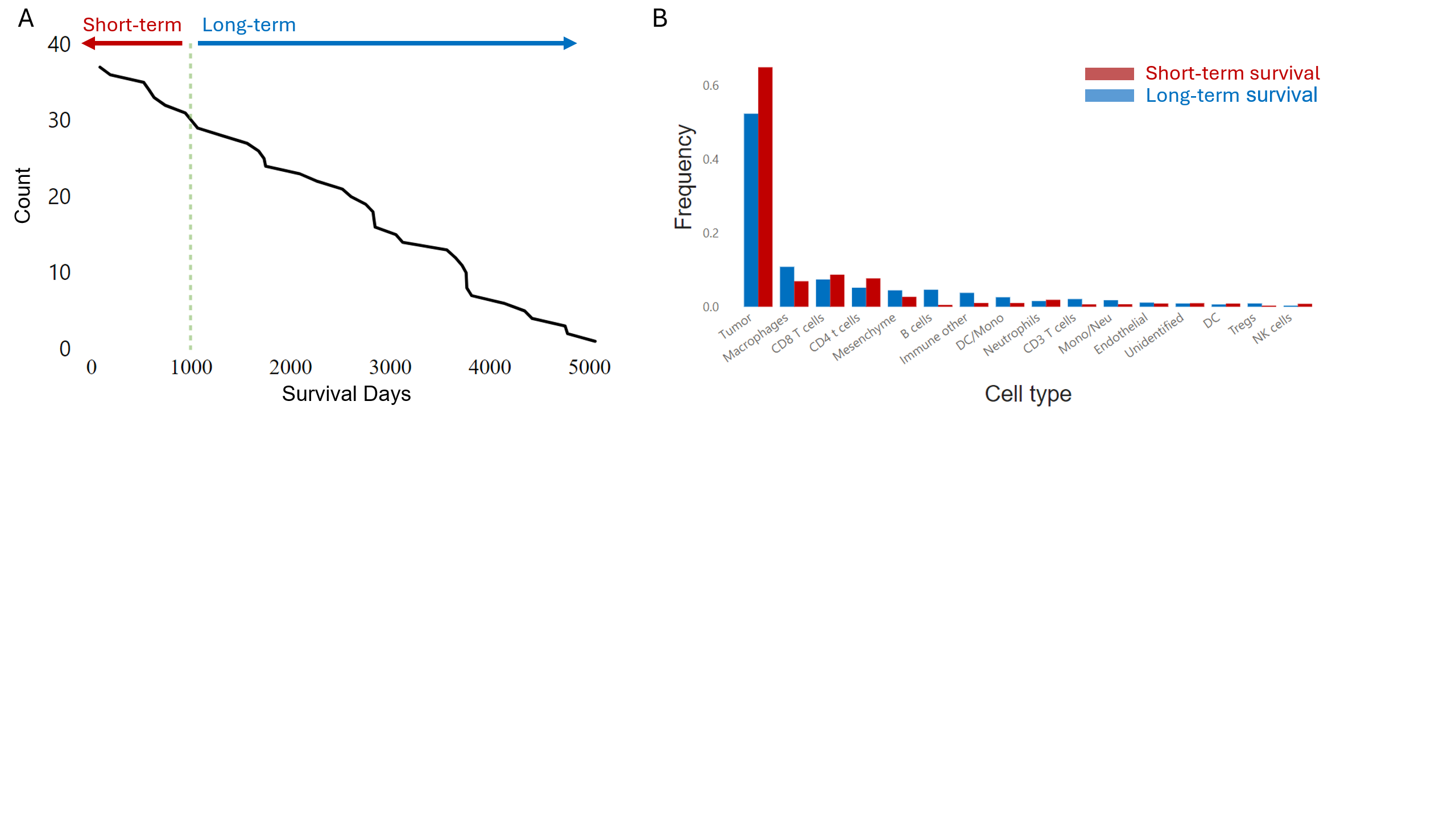
 **Figure S14. TNBC cohort.** (**A**) Number of patients who survived over time in days. The green dashed line is the cutoff between short-term (n=7) and long-term (n=30) survivors. (**B**) Cell type distribution for short-term and long-term survivors.

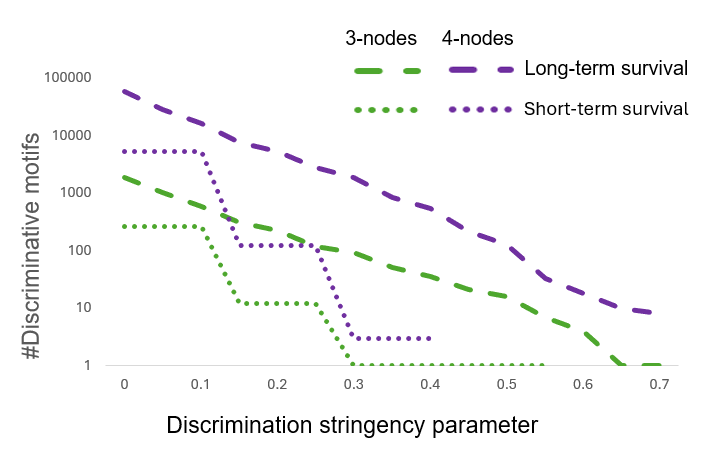

**Figure S15. Number of discriminative motifs in TNBC.** The number of 3-cell (green), 4-cell (purple) context-dependent discriminative motifs (y-axis, log scale) over the discrimination stringency parameter (x-axis).

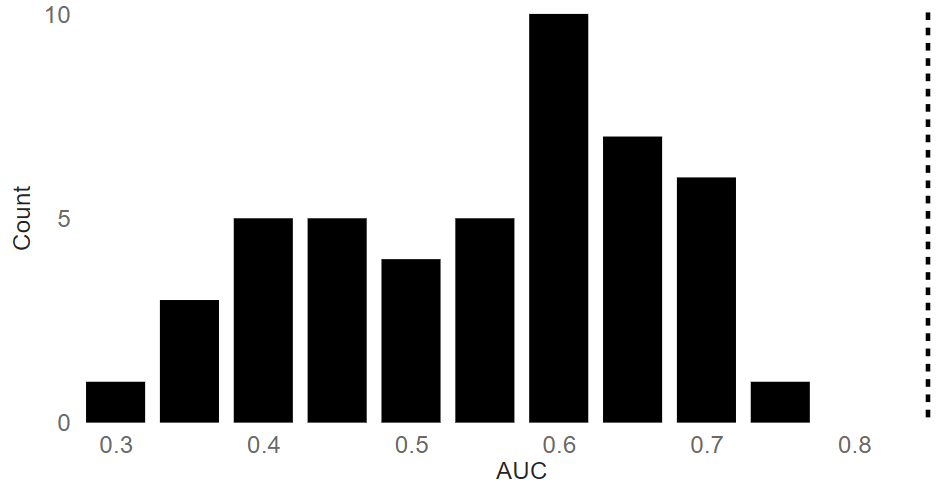

**Figure S16. Patient disease state permutation test in TNBC.** AUC histogram of 100 random permutations of the patients’ disease state. 53 permutations were excluded from the plot because they did not include discriminative features. The dashed black line is the comtrol (unpermuted) AUC result of 0.85 leading to p-value < 0.01.

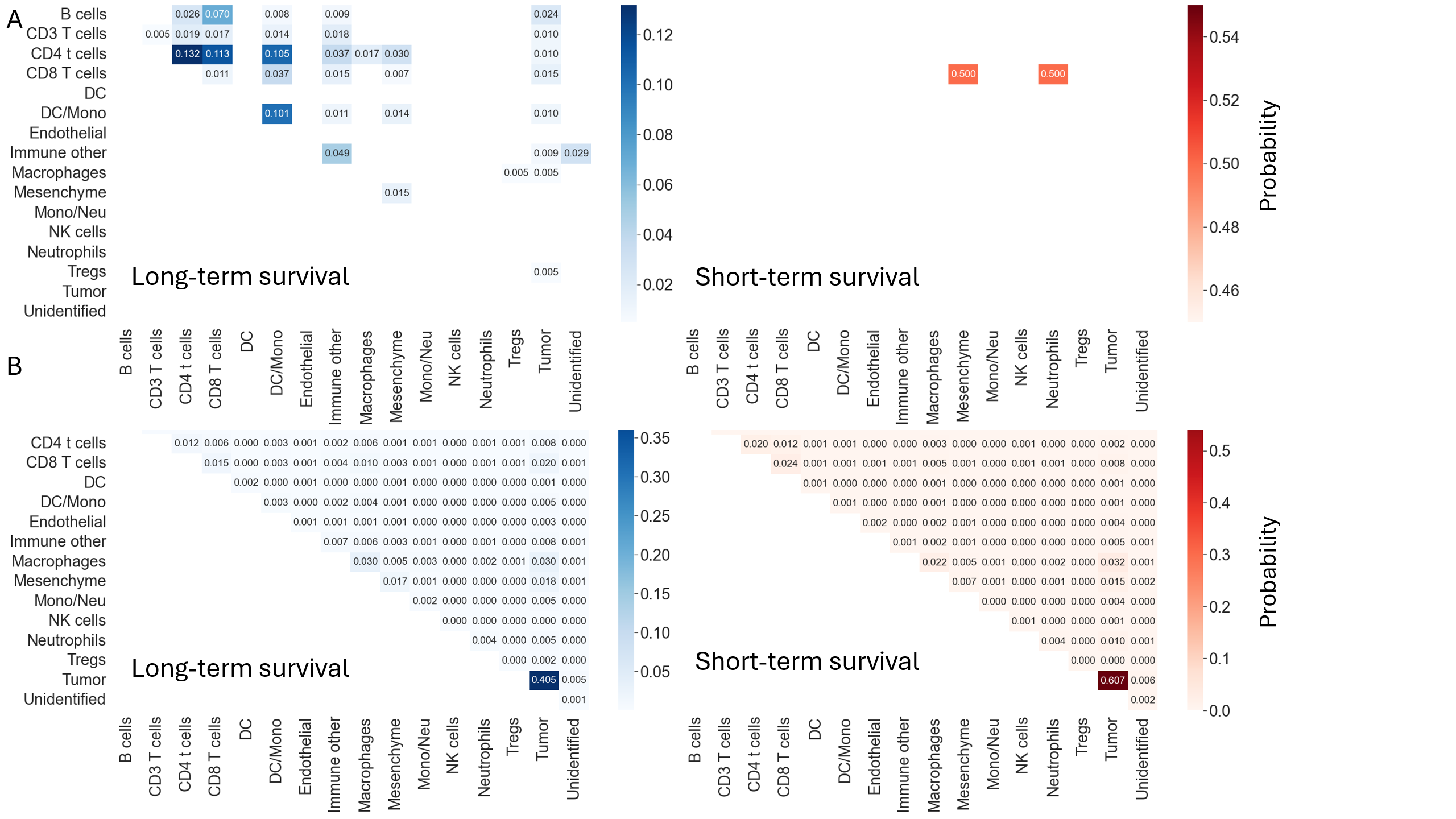
**Figure S17. TNBC discriminative motifs-induced pairwise interactions.** (**A**) The distribution of motifs-induced pairwise interactions. The probability of two cell types to be connected with an edge in the cohorts’ patients pooled set of instances of discriminative motifs associated with long-term survival patients (left**,** n = 31 motifs) and short-term survival patients (right**,** n = 1 motif). (**B**) Explicitly enumerating the distribution of pairwise interaction for long-term survival versus short-term survival patients according to the probability of two cell types being connected with an edge in the multicellular network did not show a clear distinction in the context of disease state.

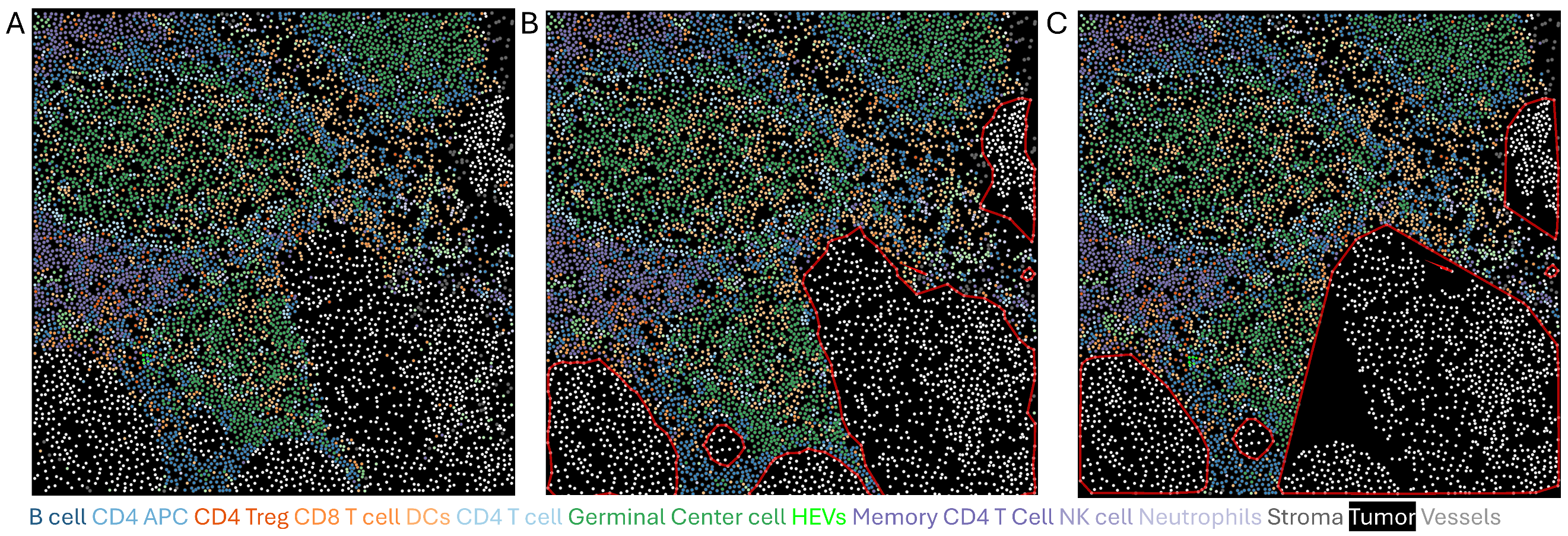

**Figure S18. Excluding tumor regions to analyze the lymph node tumor microenvironment.** An example of the tissue’s derived alpha shape for different α values. (**A**) Lymph node tissue section of patient #86 FOV #156, which includes tumor cells. **(B-C)** alpha shape clusters (red polygons) for $\alpha$ = 0.01 **(B)** versus $\alpha$ = 0 (Convex Hull)**,** creating a less accurate perimeter **(C)**. Accordingly, we set $\alpha$ to 0.01. Only tumor cells are shown inside the alpha clusters to emphasize non-tumor cell exclusion.

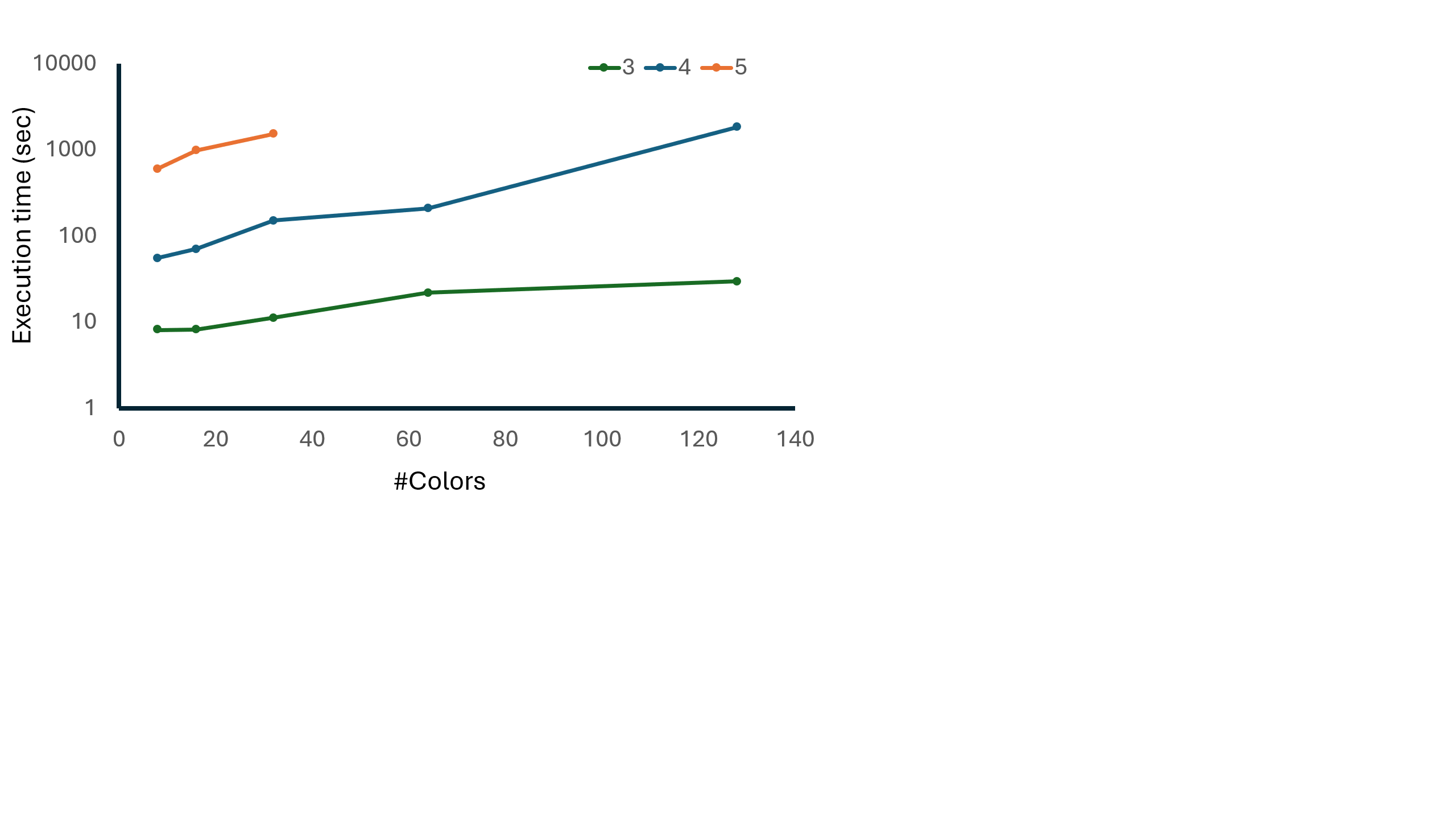

**Figure S19. FANMOD+ execution time benchmarking.** A benchmark test for the execution time (y-axis in log scale) of FANMOD+ with respect to the number of colors (x-axis) and motif size (3-nodes in green, 4-nodes in blue, and 5-nodes in orange). Each data point was calculated according to the mean execution time of five random planar graphs with n = 10,000 nodes, e ~ 30,000 edges and 8-128 colors (Methods). The calculated significance of each motif in the network (p-value) was based on 100 random networks.

### Supplementary Tables Legends

**Table S1.** Summary of the number of subgraphs and motifs of size 3-5 for each patient from the melanoma cohort. x-node-subgraphs and x-node-motifs indicate the total number of subgraphs and motifs of size x, correspondingly, across the patient’s FOVs.
